# Mechanistic Modeling and Analysis of the Mammalian Unfolded Protein Response

**DOI:** 10.1101/060020

**Authors:** Anirikh Chakrabarti, Lina Aboulmouna, Jeffrey Varner

**Affiliations:** Cornell University, School of Chemical and Biomolecular Engineering, Cornell University, Ithaca, NY 14853, USA; Purdue University, School of Chemical Engineering, West Lafayette, IN 47907, USA

## Abstract

Cells monitor protein folding by an inbuilt quality-control system in which incorrectly or misfolded folded proteins are tagged for degradation or sent back through a refolding cycle. However, continued accumulation of incorrectly folded proteins triggers the Unfolded Protein Response (UPR), which attempts to re-establish folding homeostasis or commits the cell to apoptosis. In this study, we developed a family of mechanistic models of the mammalian UPR system. An ensemble of models parameters was estimated by minimizing the difference between simulations and experimental measurements using multiobjective optimization. The ensemble of model parameters was validated using cross-validation. Analysis of the model ensemble suggested the three branches of UPR fired simultaneously. However, the importance of each brach was ranked ordered in time; PERK and IRE1 were more important early, while ATF6 was important later in the response. The activity of all three branches as coordinated by the molecular chaperone BiP. Model analysis suggested that BiP feedback was critical to the overall robustness of the system. Removal of any one branch of BiP feedback, destabilized the other branches. On the other hand, removal of all nodes of BiP feedback increased the overall robustness of the system. Thus, while BiP feedback is crucial to allowing the cell to adapt to small perturbations, it also makes the system fragile and susceptible to manipulation.

## Introduction

Protein folding is strategically important to cellular function in all organisms. In eukaryotes, secreted, membrane-bound and organelle-targeted proteins are typically processed and folded in the endoplasmic reticulum (ER)^1–3^. Intracellular perturbations caused by a variety of stressors disturb the specialized environment of the ER leading to the accumulation of misfolded or unfolded proteins ^4, 5^. Shifts in folding capacity have been associated with diseases such as cancer, diabetes and cardiovascular disorders^4^. Physiological processes such as aging can also influence protein folding. Normally, cells ensure proper protein folding using a combination of molecular chaperones, foldases and lectins^1^. However, when proper folding can not be restored, unfolded or misfolded proteins are targeted to ER Associated Degradation (ERAD) pathways for processing^3^. If unfolded or misfolded proteins continue to accumulate, eukaryotes induce the unfolded protein response (UPR). In mammalian cells, UPR is a complex signaling program mediated by three ER transmembrane receptors: activating transcription factor 6 (ATF6), inositol requiring kinase 1 (IRE1) and double-stranded RNA-activated protein kinase (PKR)-like endoplasmic reticulum kinase (PERK). UPR performs three functions, adaptation, alarm and apoptosis. During adaptation, the UPR tries to reestablish folding homeostasis by inducing the expression of chaperones that enhance protein folding. Simultaneously, translation is globally attenuated to reduce the ER folding load while the degradation of unfolded proteins is increased. If these steps fail, the UPR induces a cellular alarm and apoptosis program. The alarm phase involves several signal transduction events, ultimately leading to the removal of the translational block and the down-regulation of the expression and activity of pro-survival factors such as the B-cell lymphoma 2 (Bcl2) protein. After the alarm phase, cells can undergo apoptosis, although ER stress can also initiate autophagy^6–12^. Thus, ER folding homeostasis strongly influences mammalian physiology^5^.

ER stress and UPR plays an important role in a spectrum of diseases and technological applications^13^. In the context of diseases such as diabetes, pancreatic *β*-cells depend on efficient UPR signaling to meet the demands for constantly varying levels of insulin synthesis. Type I diabetes is marked by excessive loss of pancreatic *β*-cells, while type II diabetes is marked by pancreatic *β*-cell dysfunction. The large biosynthetic load placed on the ER by insulin production in response to blood glucose levels, can overwhelm the folding capacity of the ER. This leads to PERK activation and the reduction of protein synthesis. In PERK -/- cells, protein synthesis is unresponsive to the stress, leading to the accumulation of unfolded proteins (e.g. proinsulin) and ultimately cell death. PERK deficient mice are more prone to diabetes and progressive hyperglycemia^14^. In type II diabetes, ER stress leads to JNK-mediated phosphorylation of insulin receptor substrate 1 (IRS1) at S307, inhibiting insulin action^15^. Nitric oxide (NO), is also a key player in *β*-cell death in type-I diabetes and vascular complications in type-II diabetes. NO depletes ER Ca^2+^ leading to ER stress and ultimately apoptosis. Pancreatic *β*-cells have shown that NO-induced apoptosis is CHOP dependent^16^. Thus, ER stress is a critical feature of both type-I and type-II diabetes at the molecular, cellular and organismal level. ER stress and UPR function can also be important in cancers. The ER not only acts as the center for maturation of proteins, but also as a critical node for oxygen sensing and signaling. In rapidly growing tumors, cells face stressors like hypoxia and nutrient deprivation both of which can lead to ER stress and ultimately UPR^17^. Interestingly, the connection between hypoxia and UPR is through hypoxia inducible factor (HIF) independent pathways. Hypoxia drives PERK activation and the transient phosphorylation of eIF2*α*^18–20^on a time scale of minutes for anoxia, and much slower for hypoxic conditions^20^. Because of their ability to transduce pro-apoptotic signals, both PERK and IRE1 have been explored as potential anti-cancer targets. Versipelostatin, a repressor of BiP expression, has been shown to produce anti-tumor activity in MKN-74 xenograft mouse models^21^. Enhanced apoptosis has also been observed in BiP-deficient fibrosarcoma cells, and XBP1-and PERK-deficient mouse fibroblasts^22–24^. In the context of biotechnology, mammalian hosts, such as Chinese Hamster Ovary (CHO) cells, have been used for therapeutic protein production since the mid-1980s^25, 26^. Several current biologics are secreted, for example, the monoclonal antibodies (MAb) interferon-*γ* (IFN*γ*) or erythropoeitin (EPO)^27^. Thus, ER processing and the unfolded protein response are critical to the production of these and many other therapeutic proteins. However, despite its importance to both disease and biotechnology, there has been a paucity of mathematical models of UPR in mammalian systems.

In this study, we developed a population of mathematical models describing the adaptation, alarm, and apoptosis phases of the mammalian unfolded protein response. The biological connectivity of the UPR model was assembled following an extensive literature review. The dynamics of UPR were modeled using mass action kinetics within the framework of ordinary differential equations. An ensemble of models parameters was estimated by minimizing the difference between simulations and experimental measurements using multiobjective optimization. The ensemble of model parameters was validated using cross-validation. Analysis of the model ensemble suggested the three branches of UPR fired simultaneously. However, the importance of each brach as ranked ordered in time; PERK and IRE1 were more important early, while ATF6 was important later in the response. The activity of all three branches as coordinated by the molecular chaperone BiP. Model analysis suggested that BiP feedback was critical to the overall robustness of the system. Removal of any one branch of BiP feedback, destabilized the other branches. On the other hand, removal of all nodes of BiP feedback increased the overall robustness of the system. Thus, while BiP feedback is crucial to allowing the cell to adapt to small perturbations, it also makes the system fragile and susceptible to manipulations. The UPR model code and parameter ensemble is available under an MIT software license and can be downloaded from http://www.varnerlab.org.

## Results

### Formulation of the UPR network architecture

The UPR network described the ER folding cycle, ER-associated degradation (ERAD), ER-stress transducer (PERK, IRE1 and ATF6) signaling cascades and stress-induced caspase activation (Fig. 1). The network consisted of 636 protein or mRNA species interconnected by 1090 interactions (Fig. 1 Inset). Connectivity was formulated from a comprehensive review of the primary literature^1–5,28–34^, and from on-line databases; String-8^35^, NetworKIN^36^ and TRANSFAC. Model connectivity was not specific to a single cell-line; rather, it was a canonical representation of the pathways involved in monitoring and controlling the folding capacity of a generic well-mixed ER compartment. UPR induction was modeled as the release of BiP from the ER stress transducers, PERK, IRE1*α*, and ATF6 leading initially to adaptation of the folding cycle and then, subsequently, to alarm and apoptosis. The adaption phase of UPR was marked by general translation attenuation, selective transcriptional programs for key species like bZIP transcription factor ATF4^37^, cellular inhibitor of apoptosis (cIAP)^38^, molecular chaperones e.g., BiP^39^, and enhanced clearance of accumulated proteins via ERAD. The alarm and apoptosis phases were mediated by the induction of CHOP^40^, regulation of Bcl2, Bcl2-antagonist of cell death (BAD)^41^ and (TNF) receptor associated factor 2 (TRAF2)^34,42–44^activation (Fig. 1). Model connectivity is available from the GitHub model repository, and is described further in the supplementary materials.

**Figure 1.**
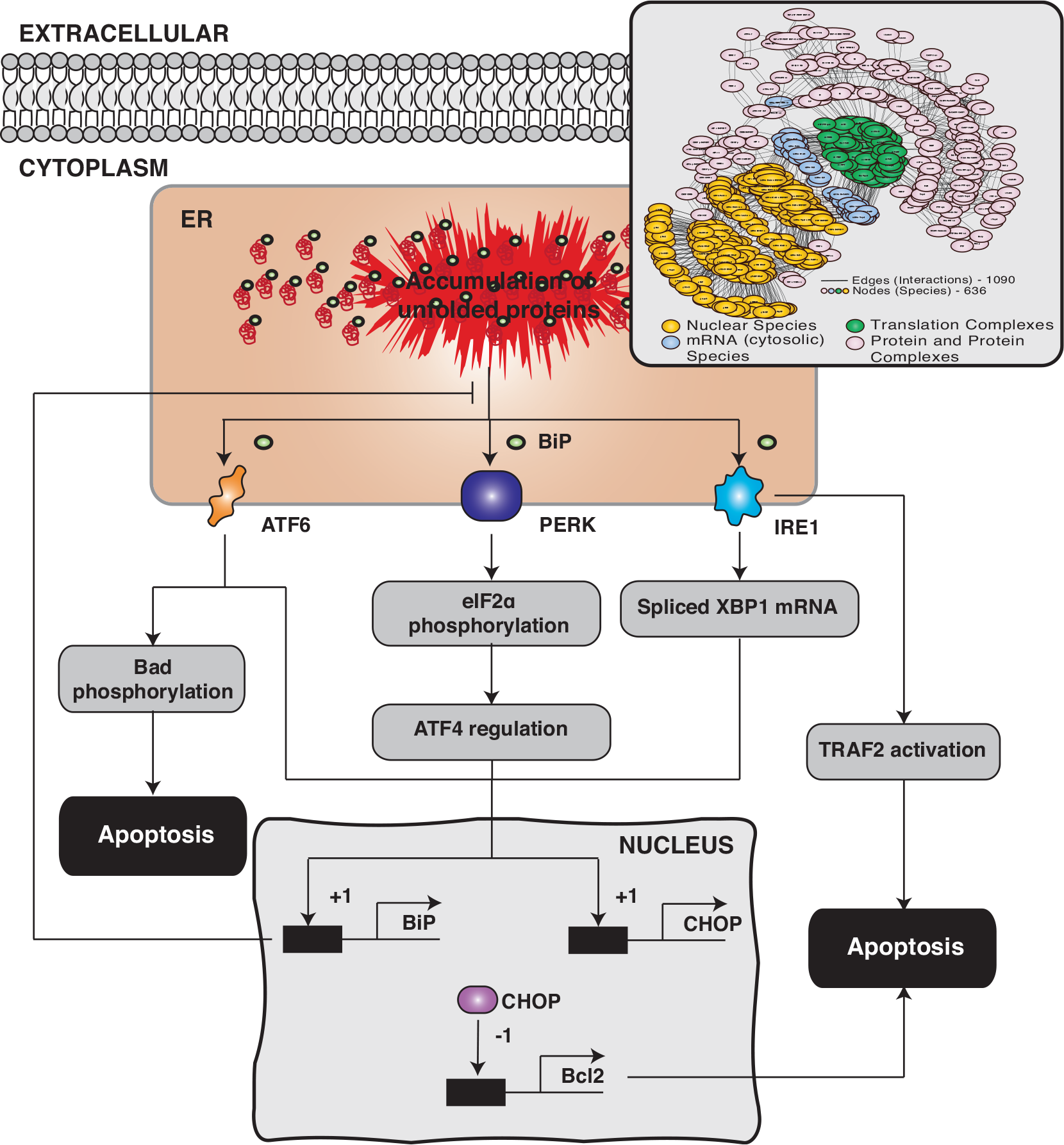
An array of cellular stressors can perturb the folding environment in the endoplasmic reticulum (ER) leading to unfolded or misfolded protein. In response to the folding imbalance, cells initiate the cytoprotective unfolded protein response (UPR). The problem of unfolded or misfolded proteins in the ER is addressed by increasing the folding capacity through the up-regulation of the expression of chaperone proteins, attenuating translation by regulating eIF2*α*, and promoting the degradation of misfolded proteins through ER-associated degradation (ERAD). If UPR is unable to restore the folding balance, ER stress will eventually lead to apoptotic cell-death. The three signal transduction pathways mediating the unfolded protein response in higher eukaryotes. First, the PRKR-like ER kinase (PERK) pathway is initiated after BiP dissociation from PERK. While PERK transduces both pro-and anti-apoptotic signals, its main function is translation attenuation through the phosphorylation of eIF2*α*. Next, the activating transcription factor 6 (ATF6) pathway is activated following BiP dissociation. ATF6 induces the expression of chaperones e.g., BiP as well as apoptosis effectors such as CHOP. Lastly, the inositol-requiring kinase 1 (IRE1) pathway is activated following BiP dissociation from IRE1. Activated IRE1 has both an endoribonuclease and a serine-threonine kinase activity that drive can pro-apoptotic signals. Inset: The UPR network consisted of 636 protein or mRNA species interconnected by 1090 interactions.

### The UPR model recapitulated the adaptation, alarm, and apoptotic phases of UPR

A population of unknown model parameters was estimated from 33 dynamic and steady state data sets taken from literature (Table T1). The residual between model simulations and each of the measurement sets was simultaneously minimized using the multiobjective POETs algorithm^45^. A leave-eight-out cross-validation strategy was used to independently estimate the training and prediction error over the 33 data sets; we estimated four different model families, where eight of the 33 objectives were reserved for validation and 25 were used for model training. Starting from an initial best-fit parameter set (nominal set), more than 25,000 probable models were estimated by POETs from which we selected N = 100 models (25 from each training family) with a Pareto rank of one or less (from approximately 1200 possible choices). The nominal, training (75 models), and prediction (25 models) errors were calculated for each objective (Table T1). Models used for prediction error calculations for a particular objective were *not* trained on that objective. The prediction likelihood was statistically significantly better for 31 of the 33 objective functions at a 95% confidence level, compared with random parameter sets generated from the nominal set (Table T1). POETs generated model families that predicted approximately 94% of the training data with a significantly higher likelihood than a random control. However, the specific value of any given parameter was likely not well described; the coefficient of variation (CV) for the model parameters ranged from 0.5 - 1.6, where approximately 65% of the parameters were constrained with a CV ≤ 1.0 (supplementary materials Fig. S1). The most constrained parameters involved a wide-array of functions e.g., regulation of PERK, eIF2*α*, ATF4, Calcineurin, BiP, CHOP and ATF6 signaling. However, the least constrained parameters involved JNK and apoptosis interactions. POETs identified Pareto fronts between several objectives, e.g., O13×O14, O25×O29, O11×O29, and O27×O2, in the training data (Fig. 2). Strong Pareto fronts suggested an inability to simultaneously model different aspects of the training data. However, fronts could also result from experimental artifacts, e.g., variation between cell-lines, time-scale differences, or from functional relationships in the data. Globally, adaptation and alarm phase training constraints conflicted with those involving apoptosis. For example, objectives involving caspase-7 or caspase-9 activity conflicted with phosphorylated eIF2*α* levels. Phosphorylation of eIF2*α* by activated PERK attenuates translation, which decreases the ER folding load. Thus, eIF2*α* phosphorylation is a key early adaptive event in UPR. On the other hand, caspase-9 is a stress-induced death marker activated only after UPR has failed to restore folding homeostasis. Conflicts between these early and late phase markers suggested the UPR time scale was perhaps cell-line or perturbation dependent. Next, we compared model simulations with measurements of components mediating the three phases of UPR.

**Figure 2.**
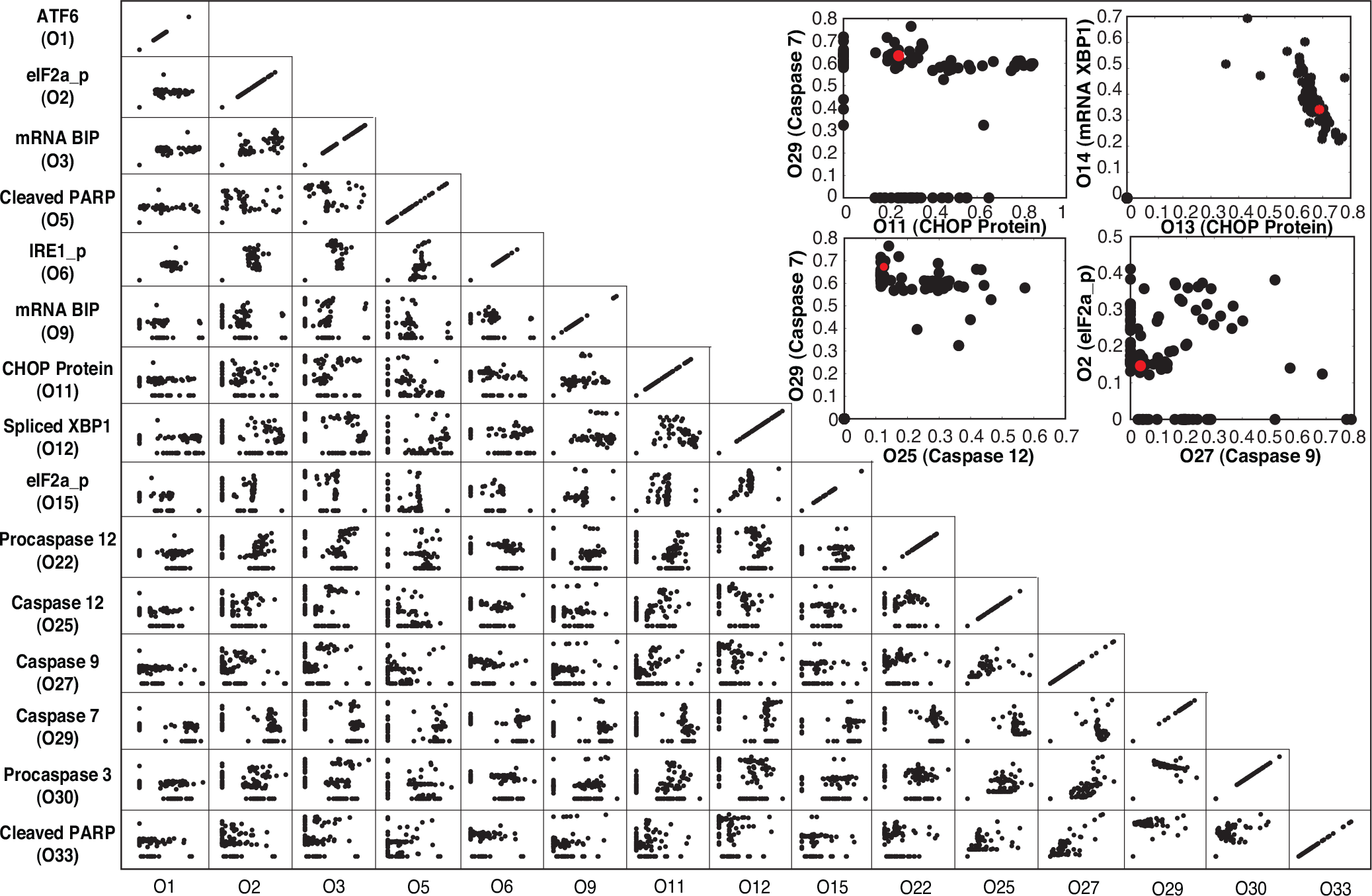
Objective function plot for selected training constrains (O1,O2,…033) for the UPR model population generated using POETs. Points denote separate models in the population. Several objectives exhibit clear Pareto fronts, e.g., O29 × O25. This suggests an inability to model both training constraints simultaneously or conflicts in the training data.

We estimated model parameters using measurements from UPR initiation and the downstream activation of apoptosis (Fig. 3). We assumed the action of thapsigargin (Tg), a non-competitive inhibitor of SERCA Ca^2+^ transporters, and other stress-inducing agents such as dithiothreitol (DTT) was identical; All stress agents led to the dissociation of BiP from the ER-stress transducers. The population of UPR models recapitulated the timescale of PERK phosphorylation (Fig. 3A) as well as its downstream signaling activity, for example, the phosphorylation of eIF2*α* (Fig. 3H). The nuclear fraction of ATF4 increased from approximately zero (untreated cells) to a maximum value 4 hrs after Tg exposure. While the model ensemble generally predicted the correct trend, there was significant error in the early time points for ATF4 (Fig. 3G). The phosphorylation of eIF2*α* by PERK is required for ATF4 activation. Interestingly, when we compared model simulations of p-eIF2*α* levels following Tg (1*μ*M) exposure in mouse embryonic fibroblasts (MEFs) with measurements (O15^46^), the model correctly captured the appropriate behavior. To test the functionality of the ATF6 branch of the UPR model, we compared simulations with measurements of cleaved ATF6 in tunicamycin-treated MEFs^47^. ER stress is known to lead to the release of BiP from ATF6. Cleaved ATF6 is then translocated to the nucleus where it up-regulates gene expression^48, 49^. Simulations of cleaved ATF6 levels following UPR initiation were consistent with measurements (Fig. 3C). Signals from the ER stress transducers converge downstream to regulate BiP transcription^50–53^. The ensemble recapitulated the correct trend BiP expression in experiments done on HEK293 cells following stress (Fig. 3D)^54^. One of the long-term outcomes of PERK/IRE1 activation is apoptotic cell death. The link between UPR and apoptosis occurred through the action of eIF2*α*, the dual role of the ATF4 transcription factor and caspase activation by the IRE1-TRAF2 signaling axis. We constrained model parameters associated with the activation of cell-death using measurements of pro/caspase-7 levels, pro/caspase-9 levels, pro/caspase-3 levels, pro/caspase-12 levels and PARP cleavage mediated by executioner caspases following treatment with 0.5*μ*M Tg^55^. These experiments were performed in Sak2 cells that lacked Apaf-1 protein expression^55^. Thus, the data allowed us to include a non-Apaf-1 mediated stress-induced caspase activation pathway into the model. The population of models recapitulated caspase-3 (Fig. 3K) and caspase-9 (Fig. 3J), as well as cleaved PARP levels (Fig. 3L) following exposure to ER stress-inducers. Interestingly, while PERK activation occurred on the timescale of minutes, initiator and executioner caspase activation occurred over 36 hrs. Thus, the population of UPR models captured complex signaling events occurring across multiple time scales. Next, we looked at the signal flow through the three branches following the induction of ER stress.

**Figure 3.**
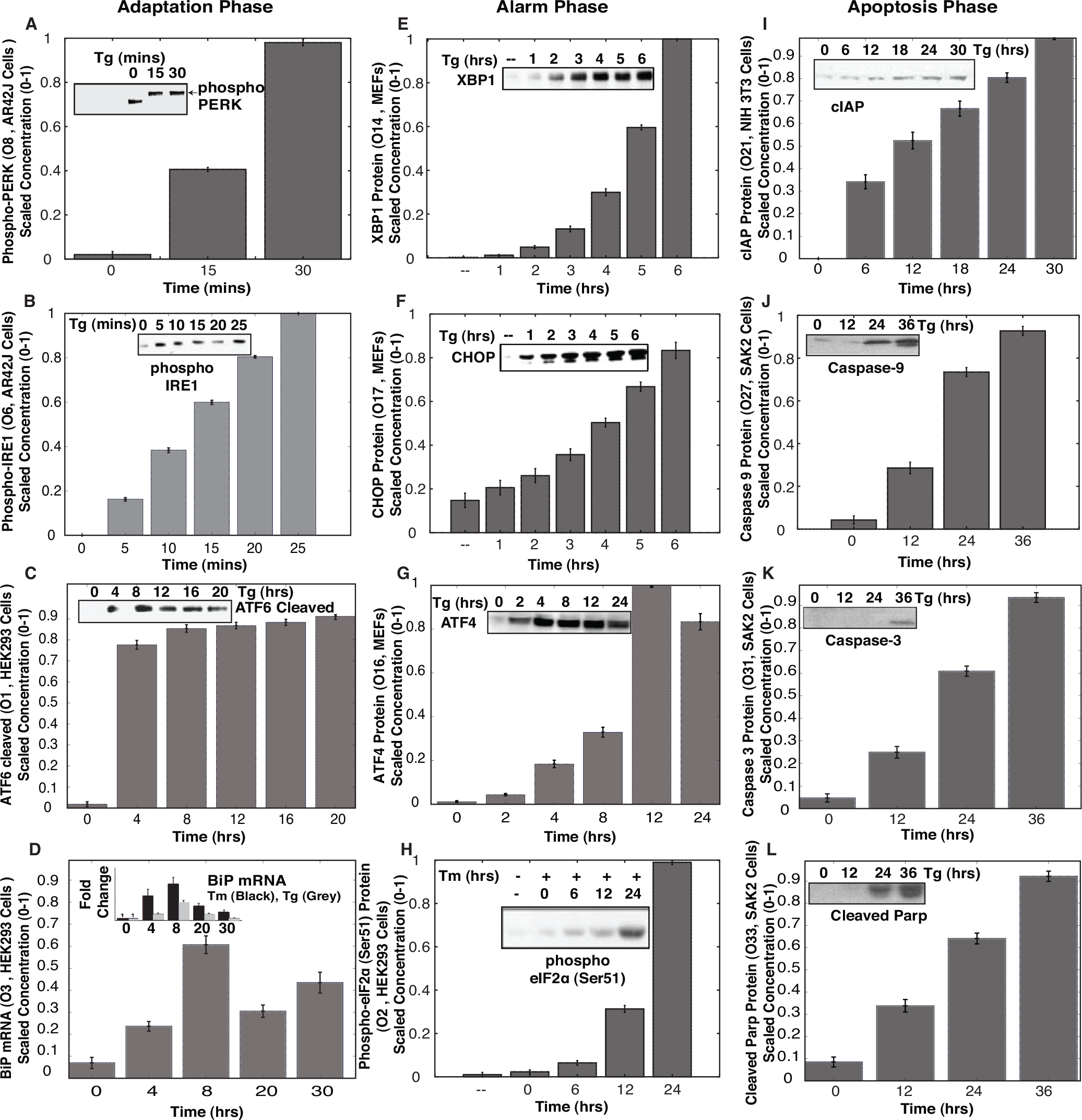
Simulations versus experimental data for selected objective functions following exposure to the ER-stress inducers Thapsigargin (Tg or Thaps) or Tunicamycin (TM). The first-column (A - D) denotes adaptation components, the second column (E - H) denotes alarm phase components, while the third column (I - L) denotes apoptosis phase components. Bars denote the scaled mean concentration computed over the ensemble, while the error bars describe one standard error.

### Signal flow analysis suggested the UPR branches fired simultaneously

Traditionally, it has been hypothesized that there is a sequential order for firing of the ER stress transducers in UPR. The PERK and ATF6 branches are thought to be activated before IRE1^34^ and largely promote ER adaptation to misfolding, while IRE1 transmits both survival and pro-apoptotic signals. However, analysis of the UPR model ensemble suggested the three branches fired simultaneously, and that adaptation, alarm, or apoptosis was the result of counteracting effects of the three UPR signaling pathways (Fig. 4 and Fig. S2). UPR induction was controlled by manipulation of the generation rate of unfolded or misfolded protein (qP) in the ER compartment. Upon induction, initially (*t* ≤ 1hr) the response was damped, marking the adaptation phase of UPR (Fig. 4A). Adaptation was followed by increased IRE1α, PERK, and ATF6 activity at *t* ~ 1 hr, marking the onset of the alarm phase. The alarm phase was followed by a steady state at *t* ~ 8-10 hrs, marking the onset of the apoptosis phase of UPR (Fig. 4A). Our analysis was substantiated further by looking at the fluxes at different phases of UPR induction (Fig. 4B). At P1, there was a marked increase in ATF4 and CHOP activity, ATF6 signaling and unfolded protein sensing and degradation by ERAD. These are hallmarks of the adaptation-alarm phase of the UPR response (Fig. 5D). The apoptosis phase at P2 (*t* ≥ 8-10 hrs) was marked by increased BiP regulation, enhanced ATF4 transcriptional activity, increased mitochondrial membrane permeability, and increased apoptotic fluxes. On the other hand, if we reduced the load of unfolded protein in the adaptation-alarm phase (P4), the cell recuperated using its ERAD machinery and the regulation of BiP (Fig. 5E-F). Thus, following initiation of UPR all three sensor branches were simultaneously activated. However, it was unclear which parameters or species controlled the responses of the three branches. We addressed this question using sensitivity and robustness analysis.

**Figure 4.**
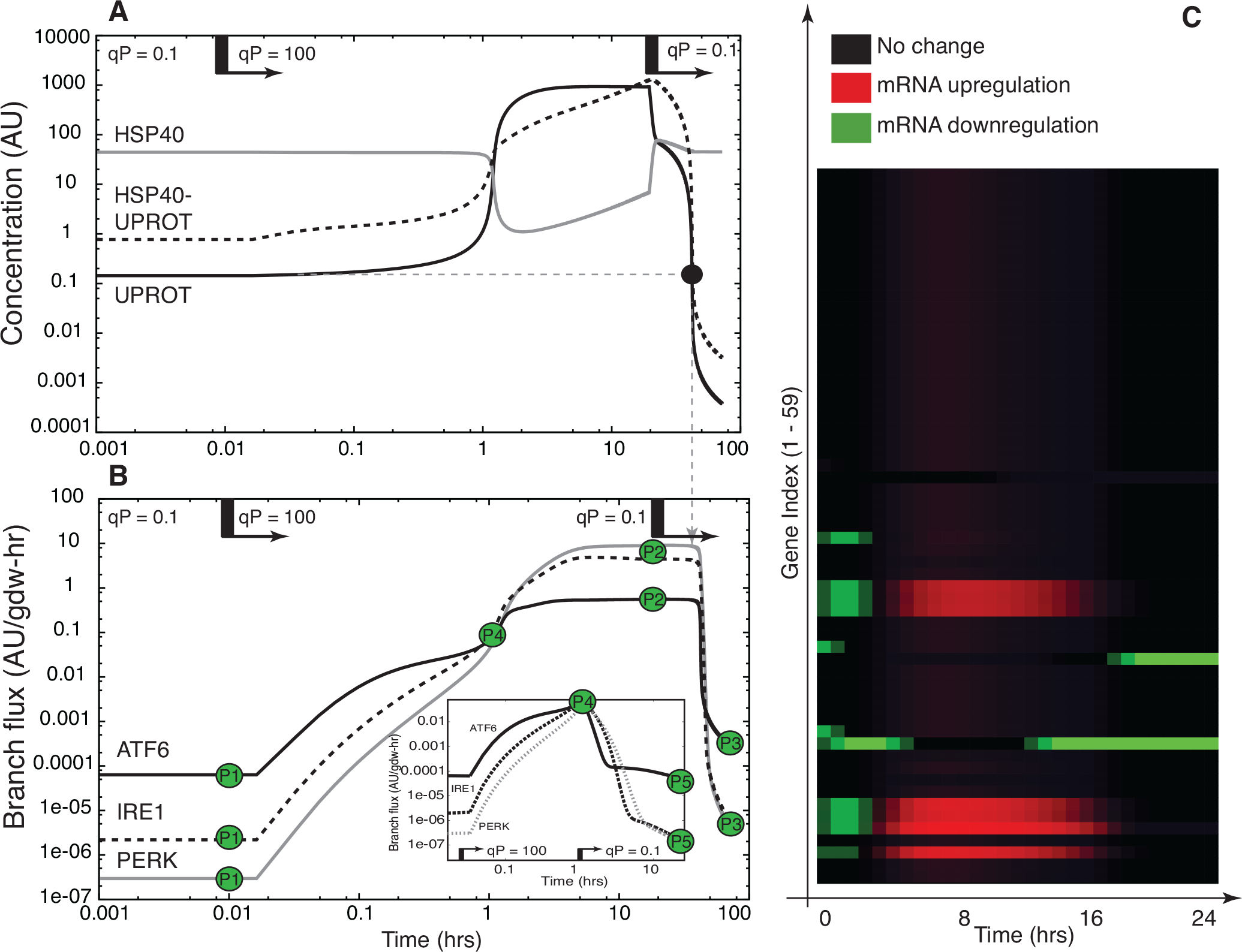
Proof of concept simulation unfolded protein response activation. **A**:UPR induction was controlled by manipulating the generation rate of unfolded or misfolded protein (qP) in the ER compartment. A step-change in qP from qP = 0.1 to qP = 100 was issued at approximately t = 0.1 hrs and then adjusted back to qP = 0.1 at t = 20 hrs. **B**:Flux through the PERK, ATF6 and IRE1 stress sensing branches as a function of time following a step change in misfolded protein generation. **C**:Simulated expression profile for the 59 genes in the model. The symbol UPROT denotes the level of unfolded protein.

**Figure 5.**
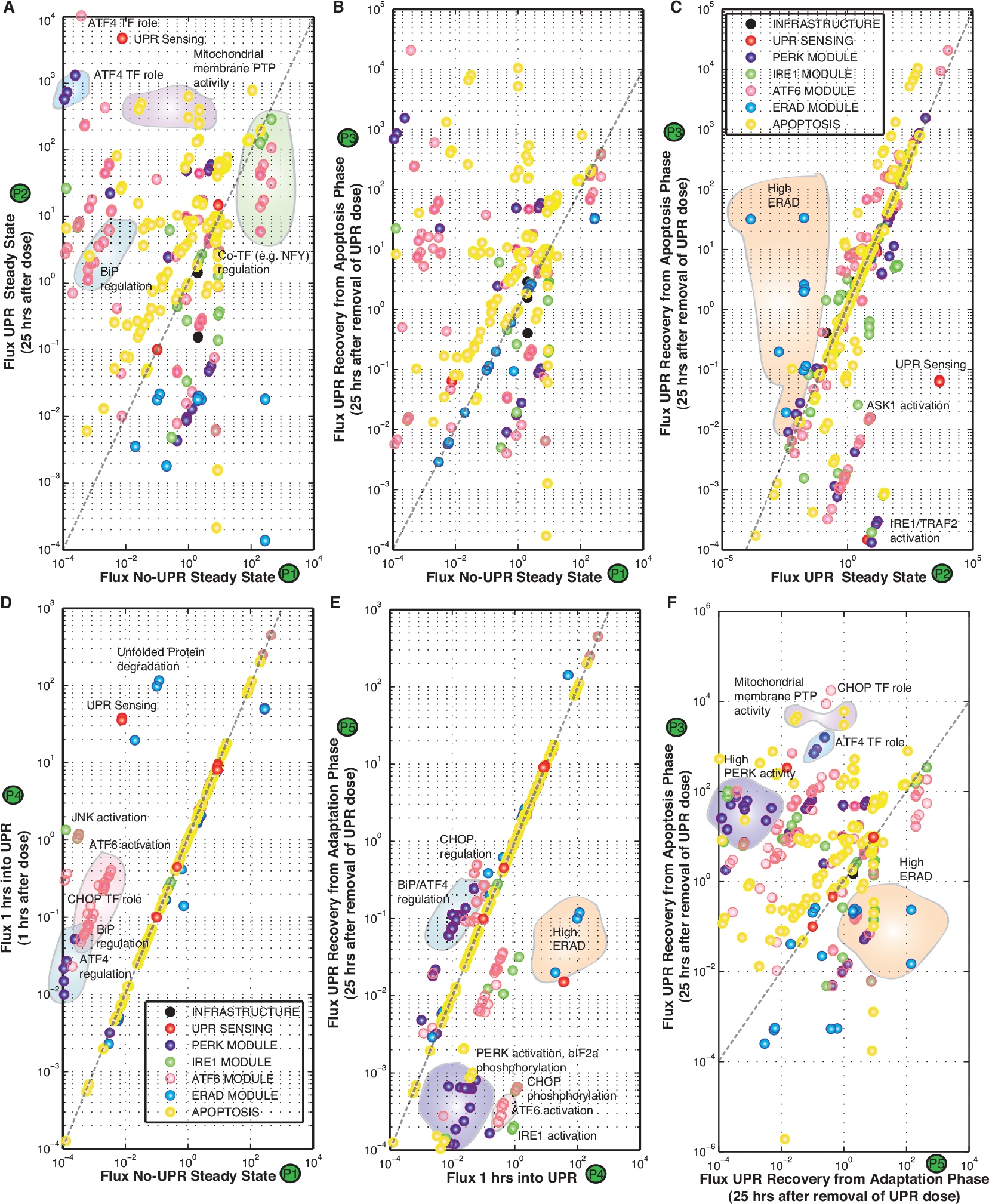
Cross plot of the fluxes at P1-P5 as denoted in Figure 4: We tried to see is how the system behaves and how the system can recuperate from UPR dose when it is in the adaptation phase as compared to the apoptosis phase. (D) As compared to P1 (No-UPR Steady State), we see that early on at 1 hr after UPR dose there is a marked increase in ATF4 and CHOP regulation, ATF6 signaling along with unfolded protein sensing and degradation. These are hallmarks of the adaptation-alarm phase of the UPR response. (A) If we continue with the dose of UPR till around 25 hrs, we see the fluxes reach a steady state. This state is marked by increased BiP regulation, enhanced ATF4 transcriptional activity, increased mitochondrial membrane permeability and increased apoptotic fluxes. This state is similar to the Apoptotic phase of UPR, where in the cell has committed itself to apoptosis mediated cell death. (B) and (C) If we reduce the UPR load after the cell has committed to apoptosis (as in P3), we find that the cell continues to function similar to the UPR state even upon UPR load reduction after 25 hrs. There are certain aspects which are seen to reduce like IRE1-TRAF2 signaling, ASK1 activation. However not much difference is seen in terms of apoptotic fluxes, denoting the cell has committed itself to death and is in a point of no return. (E)-(F) On the contrary if we reduce the load of UPR in the adaptation-alarm phase (P4), we see that the cell can recuperate using its ERAD machinery and the regulation of BiP.

First-order sensitivity coefficients were computed and time-averaged for normal and UPR induced conditions (Fig. 6). Infrastructure parameters e.g. nuclear transport, RNA polymerase or ribosome binding were globally critical, independent of stress (Fig. 6, black points). Additionally, apoptotic species and parameters were also important, both in the presence and absence of UPR (Fig. 6, yellow points). While the majority of parameters and species became more important in the presence of stress, we found a band of parameters (Fig. 6, inset) that were differentially important under stressed. The importance of the sensor and stress-transducer modules clearly increased in the presence of UPR; approximately 15% of the parameters were significantly more important in the presence of stress. These parameters were largely associated with adaptation and processing of unfolded or misfolded proteins, e.g., unfolded protein degradation, cleaved ATF6-induced gene expression, IRE1-TRAF2 mediated apoptosis regulation, and RCAN1 regulation. Sensitivity analysis conducted over discrete two hour time windows revealed the time evolution of the importance of UPR network modules (Fig. 6). Comparison of the 0 - 2 hrs time window with itself (top panel, first column of Fig. 6), supported the earlier results that infrastructure components were globally critical followed by ERAD species. These species remained important in all time windows. On the other hand, during the initial 0 - 2 hrs window, ER stress transduction pathway components were robust. Comparison of the 0 - 2 hrs time window with later time points (working down the first column, Fig. 6), showed the increasing importance of different modules as a function of time. For example, components of the PERK and IRE1 modules were more important in the 2 - 4 hrs window compared to the earlier time points, while alarm and apoptotic phase species were more important in the 6 - 8 hrs window compared to the earlier time points. Specifically, signal integration via the transcriptional activity of ATF6, ATF4, and XBP1s along with the role of RCAN1 and cIAP in apoptosis were significantly more important at 6-8 hrs as compared to 0-2 hrs time window. This was consistent with the dominant role of the negative feedback via the transcriptional regulation of BiP in UPR. Interestingly, the majority of species rankings were similar after 6 hrs (bottom row, Fig. 6 and supplementary materials Fig. S3). To further investigate the role of BiP regulation, we conducted sensitivity analysis upon knocking out the feedback branches of BiP for the nominal parameter set (Fig. S4). System performance was robust to any single knockout, however the sensitivity of the alternate feedback branches increased. The was most evident following the deletion of the ATF4 feedback. However, sensitivity coefficients are only a local measure of how small changes in parameters affect model performance. To better understand the response of UPR to a perturbation, we performed robustness analysis.

**Figure 6.**
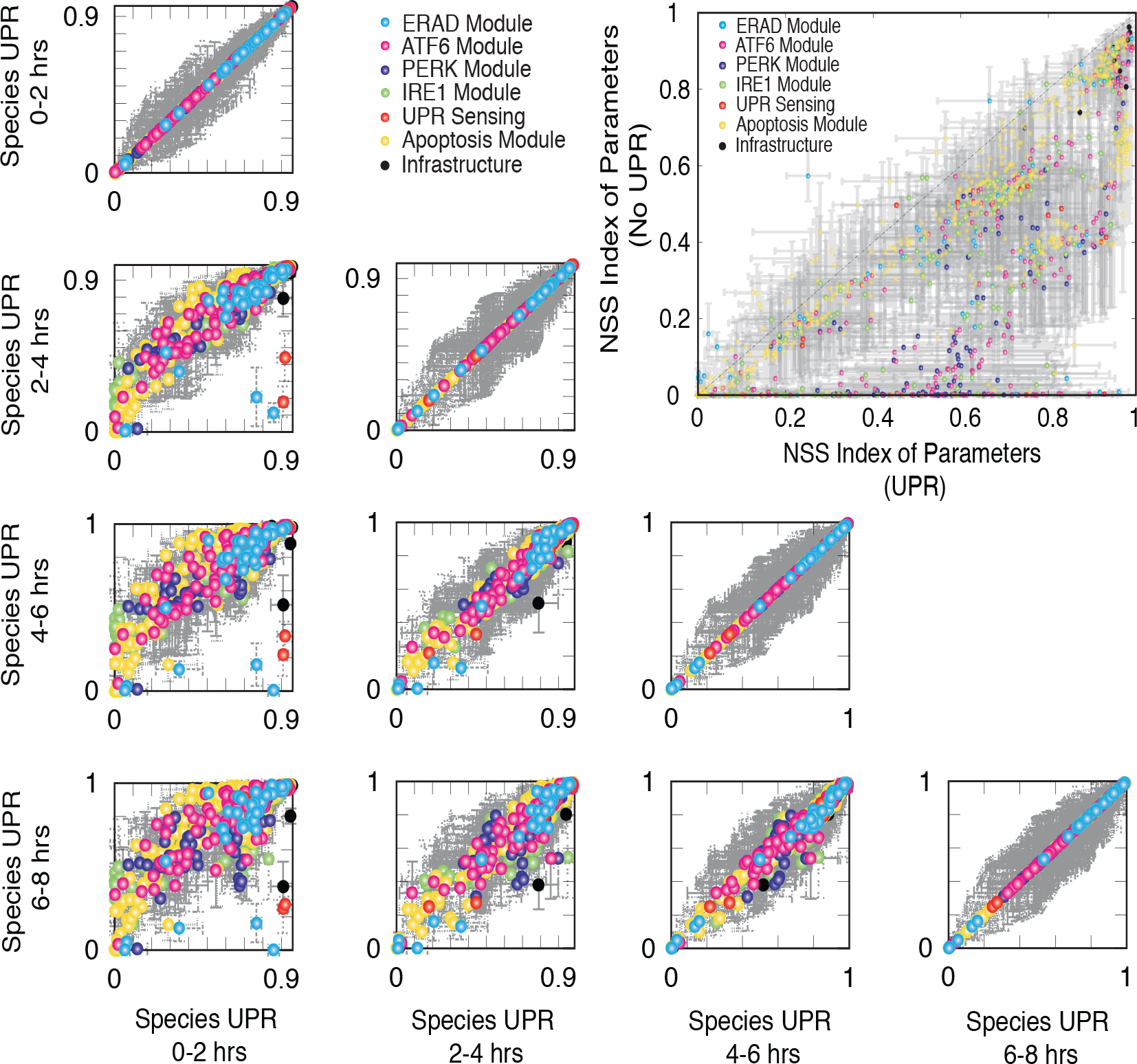
Rank-ordering of species sensitivities in the presence of UPR as a function of time. **Inset:** Rank-ordering of parameter sensitivity for UPR-induced versus normal conditions. Points denote the mean ranking computed over N = 5 parameter sets from the model population, while error bars denote one standard deviation. Points are color-coded based based upon biological function.

### Robustness analysis predicted fragile and robust structural perturbations to the UPR network

Robustness coefficients quantify the response of a protein marker to a macroscopic structural or operational perturbation to a biochemical network. We quantified shifts for 636 markers following single parameter knockouts (edge KO), single gene knockouts (GKO), and single gene overexpression (GOX) in the presence of ER stress (Fig. 7). Coefficients with values > 1 (< 1) indicated a marker increased (decreased) compared to the basal state, while a value ~ 1 indicated approximately no change following a perturbation. The cell death program (marked by robustness coefficients for caspase 3) was robust; few perturbations increased caspase 3 levels, e.g., overexpression of Procaspases 9/3 (Fig 7 A, Table T2). The robustness of caspase 3, follwed from the redundant sources of cell death (e.g., APAF-1 dependent and independent pathways). To confirm this, we simulated APAF-1 KOs over the entire ensemble (Fig. S6). We found two populations of cells in the ensemble: population 1 where APAF-1 was the dominant regulator of cell-death (marked by reduction in caspase 3 upon APAF-1 KO), and population 2 where APAF-1 was not dominant regulator. This behavior was consistent with the training data from Sak2 cells (APAF-1−ve cells)^55^. Interestingly, manipulation of the pro-survival axis via regulation of Bcl2 was possible (Fig 7A, Table T2). For example, deletion of the PERK/ATF4 signaling axes increased Bcl2 levels by removing CHOP repression (Fig. 7A). The direct correlation between ATF4 and CHOP was further observed in the ATF4-CHOP phenotypic plane. Perturbations affecting ATF4 affected CHOP levels in the same manner (Fig. 7A-B). However, owing to redundant sources of CHOP regulation (e.g., via XBP1s), effect on CHOP was damped in relation to significant changes in ATF4 levels. In the XBP1s-CHOP plane, we see at lower levels of XBP1s and CHOP, there is a direct relation between XBP1s levels and CHOP levels. However, there exists very few strategies of having both high XBP1s levels and CHOP levels indicating that higher XBP1s doesn’t necessarily mean higher CHOP levels. To further investigate the implications of the feedback regulation of BiP via ATF4/ATF6/XBP1s, we simulated KOs of these components over the entire ensemble (data not shown). Upon knockout of BiP feedback, BiP regulation was found to be very strong resulting in drastic reductions in BiP levels and ultimately a stronger and faster UPR response (Fig. S7).

**Figure 7.**
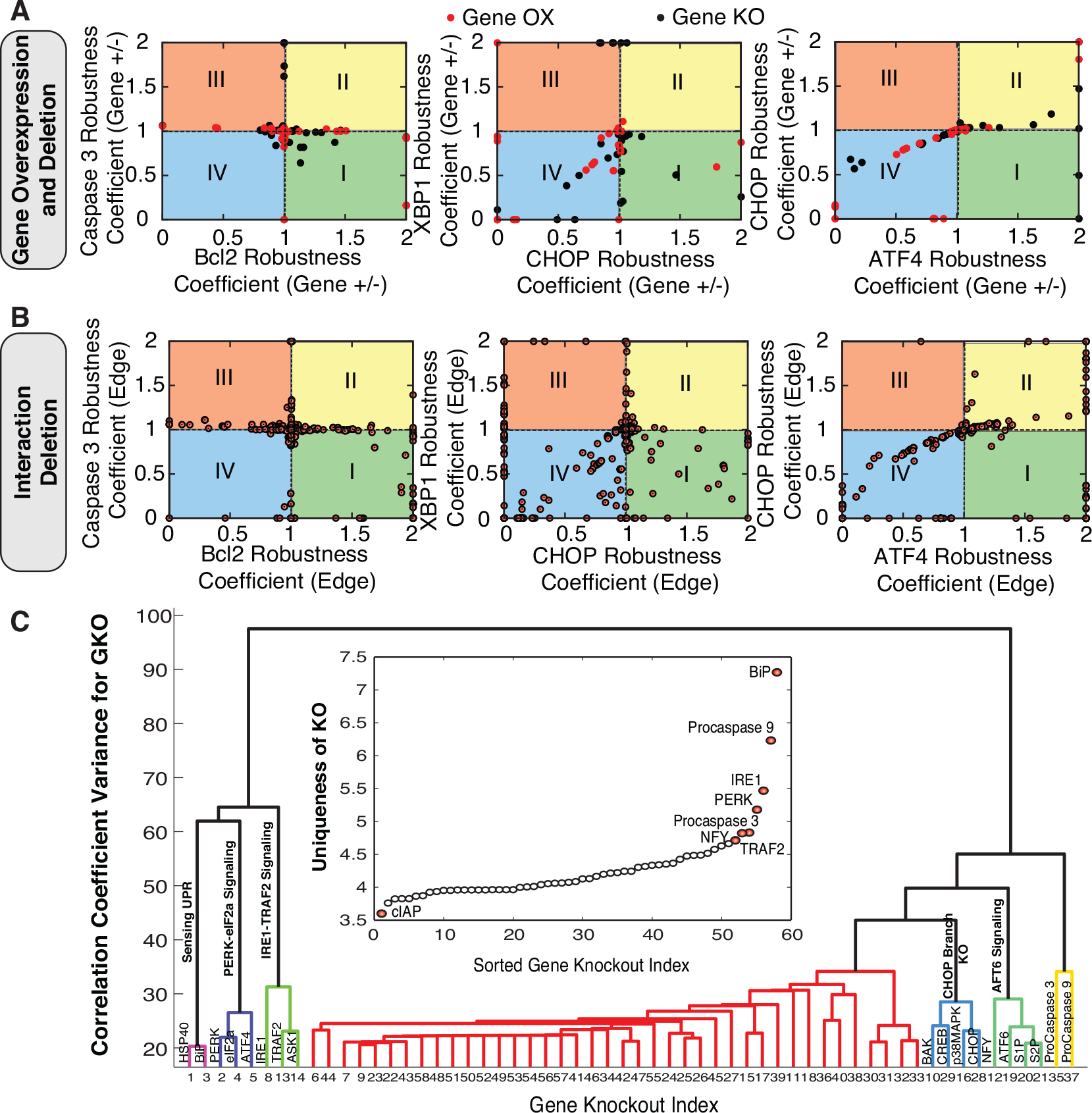
Robustness analysis of the UPR network: A-B Phenotypic phase plane analysis for the the UPR model following structural perturbations. Coupling coefficients (area under the curve from the simulation with species removed dived by the wild-type simulation) for all 636 model species were calculated for the nominal parameter set following gene overexpression/knockout (A) and deletion of single network edges (B). Coupling coefficients of one indicate no change in a marker level following a perturbation, while values less (greater) than one denote decreased (increases) marker levels. C Structural distinguishability analysis: We computed the dendrogram of the coupling coefficients for single GKO of model species. Individual coupling coefficients were clustered, where the euclidean norm was used as the distance metric and the linkage function was the inner square product (variance minimization algorithm). Each additional cluster was chosen to reduce the overall variance (y-axis). A general description of the biological function of the clusters were indicated by each group. **Insets:** Distinguishability as the magnitude of the orthogonal components for all knockout species. Species were ordered from largest to smallest magnitudes. Red markers indicate species which were statistically significant.

Clustering of the gene knockout robustness coefficients provided systems-level insight into UPR (Fig. 7C). The most distinct separation was between the ATF6 and IRE1/PERK branches. PERK/ATF4 plays a dominant role in the regulation of BiP and CHOP upon the onset of UPR. Similarly, the IRE1/TRAF2 signaling axis induced apoptosis. Another interesting functional module was that of CHOP, involving p38MAPK which leads to down-regulation of Bcl2 levels which considerably affects the apoptosis module. We computed the magnitude of the orthogonal components, which was used to establish the uniqueness of a single gene knockout (Fig. 7C Inset). BiP, Procaspase 9, IRE1, PERK and TRAF2 knockouts produced the most unique effects. This was supported by the critical role of BiP in initiating and regulating the time scale of the UPR response. PERK (via ATF4) plays a key role in the regulation of BiP, thus its uniqueness. Similarly, the regulation of the apoptosis branch via IRE1-TRAF2 and Procaspase 9 were the amongst the most unique.

## Discussion

In this study, we developed a population of mathematical models describing the adaptation, alarm, and apoptosis phases of the mammalian unfolded protein response (UPR). Proteins requiring post-translational modifications such as N-linked glycosylation or disulfide bond formation are processed in the endoplasmic reticulum (ER). A diverse array of cellular stresses can lead to dysfunction of the ER, and ultimately to an imbalance between protein-folding capacity and protein-folding load. Unfolded or misfolded proteins are tagged for degradation via ER associated degradation (ERAD) or sent back through the folding cycle. Continued accumulation of incorrectly folded proteins can also trigger UPR. PRKR-like ER kinase (PERK), inositol-requiring kinase 1 (IRE1) and activating transcription factor 6 (ATF6) were modeled as the key UPR initiators. While UPR has been extensively studied^1–5,28–34^, a detailed mathematical model of this important system has not been developed. The UPR network architecture used in the model was based on extensive review of the literature (supplementary materials). Mass balance equations describing 636 species interconnected by 1090 interactions were formulated using mass-action kinetics within an ordinary differential equation (ODE) framework. Four model populations were estimated using multi-objective optimization (33 objective functions) in conjunction with a leave-eight out cross-validation strategy using POETs^45^. These model populations were then analyzed using population-based sensitivity and robustness analysis.

A key finding of our study was that the overall outcome of UPR was as a result of simultaneous firing and competition between signaling mediated by the three ER-stress transducers: PERK, IRE1, and ATF6. This is in contrast to the traditional belief that PERK and ATF6 branches are activated before IRE1^34^. So what we hypothesize is that instead of a sequential ordering of these branches, the state of the cell in terms of adaptation, alarm, or apoptosis is a result of counteracting effects of these three prongs of UPR signaling. The counteracting/competing effects were further substantiated in simulated knockout studies, wherein knockout of one ER stress transducer led to enhancement of the other branches of UPR. Signal transduction architectures frequently contain redundancy, feedback, and crosstalk. These topological features ensure signal propagation is adaptable, efficient, and robust. However, they also make reprogramming signal flow challenging. This was highlighted remarkably in the case of UPR. Signals from the three ER-stress transducers converged at the level of up-regulation of BiP. This is the key junction which regulates the three stages (onset and time) of adaptation, alarm, and apoptosis. In this regard, regulators of this feedback cleaved ATF6, ATF4, and XBP1s and were seen as highly sensitive components of UPR. Interestingly these components put-together increased the overall fragility of the system and presented a greater scope of manipulation of the UPR response. This was substantiated by sensitivity analysis upon KO of the feedback loops, where we saw increased stability of the UPR module. When these feedback components were knocked out individually, the system overall remained stable thanks to increased activity and load sharing via the other feedback branches. Amongst the three components of feedback, we identified ATF4 as the key load bearer/regulator. This was substantiated by signal flow, robustness and sensitivity analysis. This is really interesting as ATF4 protein has shown to be present in greater levels in cancer compared to normal tissue, and it is up-regulated by signals of the tumor microenvironment such as hypoxia/anoxia, oxidative stress, and ER stress^56^. So any aberrations in regulation of ATF4 could potentially serve as a specific target in cancer therapy. As a target, ATF4 is attractive because it is also potentially involved in angiogenesis and adaptation of cancer cells to hypoxia/anoxia, which are major problems in cancer progression^56^.

Downstream effects of UPR range from cellular adaptation/survival (low stress) to the cell committing to apoptosis mediated death (high stress). Our modeling analysis suggested that the cell-death phenotype (marked by increased levels of Caspase 3 as compared to WT) was relatively robust. This robustness could be attributed to redundant routes of APAF-1 dependent and APAF-1 independent routes of apoptosis. This claim is supported by experimental evidence in Sak2 cells^55^ and as seen by our simulated knockout studies where we identified two distinct populations representing clear distinctions in APAF-1 dependent and independent routes of apoptosis. Interestingly, manipulation of the pro-survival phenotype (marked by increased levels of Bcl2 as compared to WT) was feasible. The most effective route was via manipulation of the PERK/ATF4/CHOP branch. This was substantiated by simulated CHOP KO experiments over the entire ensemble, wherein we identified two distinct populations within the ensembles. One with a strong effect of CHOP mediated down-regulation of Bcl2 (marked by ~ 10 fold increase in Bcl2 levels) and the other with very little effect of CHOP on Bcl2 levels. This complex network behavior could be attributed to other conflicting means of regulation of Bcl2 levels. Rightfully so, induction of CHOP is involved in the development of various diseases and several therapeutic interventions^57^. For instance, suppression of CHOP by RNA interference, decoy oligodeoxynucleotides or drug inhibitors have a significant therapeutic potential to modulate type I diabetes and brain ischemia. On the other hand, overexpression of CHOP may represent a new class of anticancer therapy. Since induction of BiP has been observed in a variety of tumor cells, overexpression of CHOP directed by the BiP promoter may be used as a highly specific therapy for cancer^57^. Model analysis also highlighted the essential role of RCAN1 and IRE1-TRAF2 routes of apoptosis. ATF6 induces regulation of calcineurin 1 (RCAN1) expression^31^. RCAN1 sequesters calcineurin^31^, a calcium activated protein-phosphotase B, that dephosphorylates Bcl2-antagonist of cell death (BAD) at S75 or S99^41^. This leads to sequestering of Bcl2 by BAD, which inhibits its downstream anti-apoptotic activity^41^. Recently, a number of ATF6 homologs have been identified, e.g., OASIS, CREBH, LUMAN/CREB3, CREB4, and BBF2H7 that are processed in a similar way as ATF6, yet their function remains unknown^58^. Thus, ER-stress induced ATF6 signaling may be responsible for additional undiscovered functionality.

We generated insights and presented falsifiable hypothesis regarding the UPR program using mathematical modeling. While we did an extensive literature search to formulate the model, we are likely missing key structural interactions in the UPR interaction network. First, we are missing the negative regulation of the three ER-stress transducers. Given PERK’s central role in translation attenuation, cells have evolved multiple mechanisms to regulate PERK activity^59^. The cytosolic kinase domain of PERK can be inhibited by the action of the DNAJ family member P58^*Ipk*^. P58^*Ipk*^ was initially discovered as an inhibitor of the eIF2*α* protein kinase PKR^59^. P58^*Ipk*^, whose expression is induced following ATF6 activation, binds to the cytosolic kinase domain of PERK, inhibiting its activity^60, 61^. Inhibition of PERK kinase activity relieves eIF2*α* phosphorylation, thereby removing the translational block. Interestingly, P58^*Ipk*^ expression occurs several hours after PERK activation and eIF2*α* phosphorylation. Thus, P58^*Ipk*^ induction may mark the end of UPR adaptation, and the beginning of the alarm/apoptosis phase of the response^34^. Second, PERK induces a negative feedback loop, through its downstream effector CHOP, involving the de-phosphorylation of eIF2*α*. CHOP induces the expression of GADD34 which, in conjunction with protein phosphatase 1 (PP1), assembles into a phosphatase which dephosphorylates the S51 residue of eIF2*α*^62^. GADD34 is a member of the GADD family of genes which are induced by DNA damage and a variety of other cellular stresses^63^. The GADD34 binding partner in this complex appears to be responsible for PP1α recognition and targeting of the phosphatase complex to the ER. Association between GADD34 and PP1 is encoded by a C-terminal canonical PP1 binding motif, KVRF, while approximately 180 residues near the N-terminus of GADD34, appear to be responsible for ER localization^64^. Next, little is known about deactivation of ATF6. Recently, XBP1u, the unspliced form of XBP1, has been implicated as a negative regulator for ATF6^65^. In the recovery phase following ER stress, high levels of XBP1u may play a dual role by promoting degradation^66,67^, and binding of ATF6*α* rendering it prone to proteasomal degradation^65^. Lastly, we should revisit the regulation of IRE1*α* activity. IRE1*α* activity is regulated by several proteins, including tyrosine phosphatase 1B (PTP-1B), ASK1-interactive protein 1 (AIP1) and members of the Bcl2 protein family. Members of the HSP family of proteins have also been shown to regulate IRE1*α*. For example, HSP90 interacts with the cytosolic domain of IRE1*α*, potentially protecting it from degradation by the proteasome^68^. HSP72 interaction with the cytosolic IRE1*α* domain has also recently been shown to enhance IRE1*α* endoribonuclease activity^69^. These missing structural connections could be important to fully understanding and manipulating UPR.

## Methods

### Formulation and solution of the model equations

The unfolded protein response model was formulated as a set of coupled ordinary differential equations (ODEs):

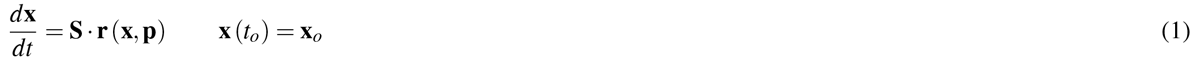

The symbol S denotes the stoichiometric matrix (636 × 1090). The quantity x denotes the concentration vector of proteins or protein complexes (636 × 1). The term **r** (**x**, **p**) denotes the vector of reaction rates (1090 × 1). Each row in **S** described a protein or protein-protein complex, while each column described the stoichiometry of network interactions. Thus, the (*i*, *j*) element of **S**, denoted by ***σ***_*ij*_, described how protein *i* was involved in rate *j*. If ***σ***_*ij*_ < 0, then protein *i* was consumed in *r_j_*. Conversely, if ***σ***_*ij*_ > 0, protein *i* was produced by *r_j_*. Lastly, if ***σ***_*ij*_ = 0, there was no protein *i* in rate *j*. All of these interactions were obtained from the literature (supplemental materials). We assumed mass-action kinetics for each interaction in the network. The rate expression for interaction *q* was given by:

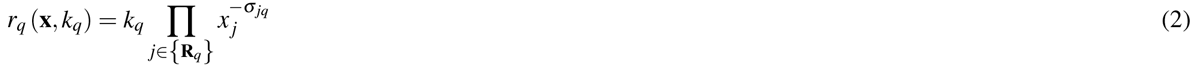

The set {**R**_*q*_} denotes reactants for reaction *q* while ***σ***_*jq*_ denotes the stoichiometric coefficient (element of the matrix **S**) governing species *j* in reaction *q*. All reversible interactions were split into two irreversible steps. The mass-action formulation, while expanding the dimension of the UPR model, regularized the mathematical structure. Parameters were one of only three types: association, dissociation, or catalytic rate constants. Thus, although mass-action kinetics increased the number of parameters and species, they reduced the complexity of model analysis. In this study, we considered well-mixed nuclear, cytosolic, and extracellular compartments. Unfolded protein response conditions were simulated by running the model to steady state and then providing a dose of unfolded protein. The steady-state was estimated numerically by repeatedly solving the model equations and estimating the difference between subsequent time points:

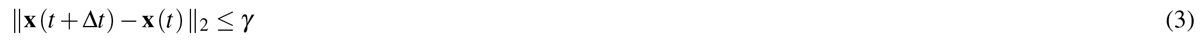

The quantities **x** (*t*) and **x** (*t* + Δ*t*) denote the simulated concentration vector at time *t* and *t* + Δ*t*, respectively. The *L*_2_ vector-norm was used as the distance metric. We used Δ*t* = 1 s and *γ* = 0.001 for all simulations. The model equations were solved using the LSODE routine in OCTAVE (http://www.octave.org) on an Apple workstation (Apple, Cupertino, CA). The UPR model code and parameter ensemble is available under an MIT software license and can be downloaded from http://www.varnerlab.org.

### Estimation and cross-validation of a population of UPR models

POETs is a multiobjective optimization strategy which integrates several local search strategies e.g., Simulated Annealing (SA) or Pattern Search (PS) with a Pareto-rank-based fitness assignment^45^. Let **k**_*i*+1_ denote a candidate parameter set at iteration *i* + 1. The squared error for **k**_*i*+1_ for training set *j* was defined as:

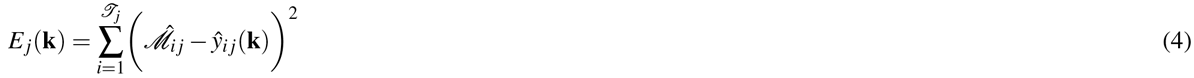

The symbol 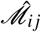 denotes scaled experimental observations (from training set *j*) while the symbol *ŷ*_*ij*_ denotes the scaled simulation output (from training set *j*). The quantity *i* denotes the sampled time-index and 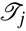 denotes the number of time points for experiment *j*. The read-out from the training immunoblots was band intensity where we assumed intensity was only loosely proportional to concentration. Suppose we have the intensity for species *x* at time *i* = {*t*_1_, *t*_2_,‥, *t_n_*} in condition *j*. The scaled measurement would then be given by:

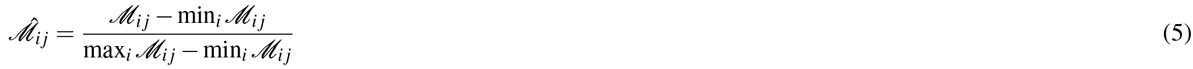

Under this scaling, the lowest intensity band equaled zero while the highest intensity band equaled one. A similar scaling was defined for the simulation output.

We computed the Pareto rank of **k**_*i*+1_ by comparing the simulation error at iteration *i* + 1 against the simulation archive **K**_*i*_. We used the Fonseca and Fleming ranking scheme^70^:

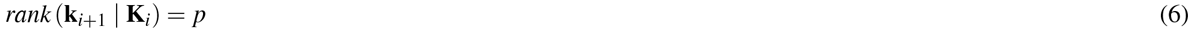

where *p* denotes the number of parameter sets that dominate parameter set **k**_*i*+1_. Parameter sets on or near the optimal trade-off surface have small rank. Sets with increasing rank are progressively further away from the optimal trade-off surface. The parameter set **k**_*i*+1_ was accepted or rejected by the SA with probability 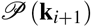:

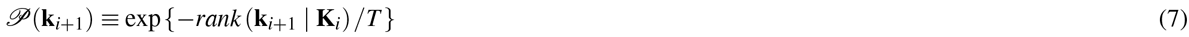

where *T* is the computational annealing temperature. The initial temperature *T_o_* = *n*/*log*(2), where *n* is user defined (*n* = 4 for this study). The final temperature was *T_f_* = 0.1. The annealing temperature was discretized into 10 quanta between *T_o_* and *T_f_* and adjusted according to the schedule *T_k_* = *β^k^T*_0_ where *β* was defined as:

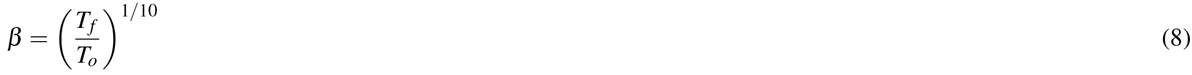

The epoch-counter *k* was incremented after the addition of 100 members to the ensemble. Thus, as the ensemble grew, the likelihood of accepting parameter sets with a large Pareto rank decreased. To generate parameter diversity, we randomly perturbed each parameter by ≤ ±25%. We performed a local pattern-search every *q* steps to minimize the residual for a single randomly selected objective. The local pattern-search algorithm has been described previously^71, 72^. The parameter ensemble used in the simulation and sensitivity studies was generated from the low-rank parameter sets in **K**_i_.

We simultaneously calculated training and prediction error during the parameter estimation procedure using leave-eight-out cross-validation^73^. The complete set of training data (33 objectives) was subdivided into four bins; in each bin 25 data sets were reserved for training while eight were reserved for prediction. In the first bin DS_1_…DS_8_ were used for validation while DS_9_…DS_33_ were used for training. In the second bin DS_9_…DS_16_ were used for validation while DS_1_…DS_8_ DS_17_…DS_33_ were used for training, etc. Thus, we formulated four ensembles from which we evenly selected parameter sets for the *parent* ensemble. While cross-validation required that we generate additional model populations, we trained and tested against all the data sets.

### Sensitivity and robustness analysis of the population of UPR models

Sensitivity coefficients were calculated as shown previously^45^ using five models selected from the ensemble (red points, supplementary materials Fig. S1). The resulting sensitivity coefficients were scaled and time-averaged (Trapezoid rule):

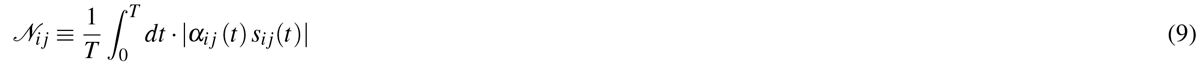

*T*
where *T* denotes the final simulation time and *α_ij_* = 1. The time-averaged sensitivity coefficients were then organized into an array for each ensemble member:

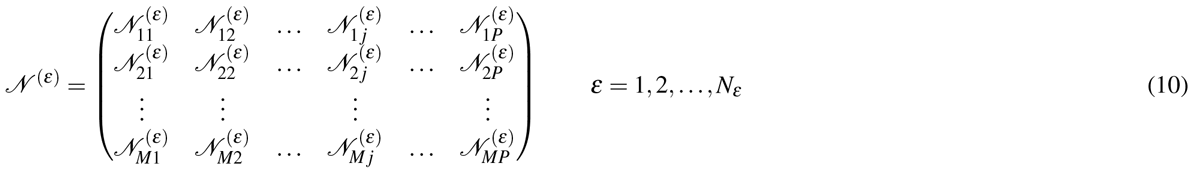

where *ɛ* denotes the index of the ensemble member, *P* denotes the number of parameters, *N_ɛ_* denotes the number of ensemble samples and *M* denotes the number of model species. To estimate the relative fragility or robustness of species and reactions in the network, we decomposed the 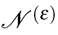 matrix using Singular Value Decomposition (SVD):

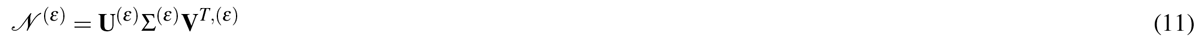

Coefficients of the left (right) singular vectors corresponding to largest *β* singular values of 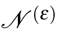 were rank-ordered to estimate important species (reaction) combinations. Only coefficients with magnitude greater than a threshold (*δ* = 0.1) were considered. The fraction of the *β* vectors in which a reaction or species index occurred was used to rank its importance.

Robustness coefficients of the form:

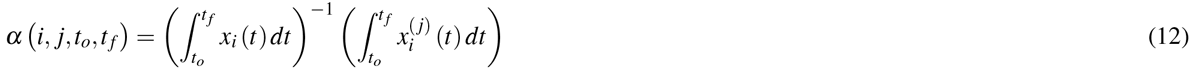

were calculated to understand the robustness of the network. The robustness coefficient *α* (*i*, *j*, *t_o_*, *t_f_*) is the ratio of the integrated concentration of a network marker in the presence (numerator) and absence (denominator) of structural or operational perturbation. The quantities *t*_0_ and *t_f_* denote the initial and final simulation time respectively, while *i* and *j* denote the indices for the marker and the perturbation respectively. If (*i*, *j*, *t_o_*, *t_f_*) > 1, then the perturbation *increased* the marker concentration. Conversely, if (*i*, *j*, *t_o_*, *t_f_*) ≪ 1 the perturbation *decreased* the marker concentration. Lastly, if (*i*, *j*, *t_o_*, *t_f_*) ~ 1 the perturbation did not influence the marker concentration.

### Clustering and identification of distinguishable species

A dendrogram was derived by considering each of the knockouts(over-expressions) as variables and the average log of robustness coefficient (LRC) for each of the species as observations. We used the Euclidean norm in LRC space as the distance metric. The linkage function (objective function for identifying variable clusters) was the inner squared distance (minimum variance algorithm). The Statistical Toolbox of Matlab (The Mathworks, Natick, MA) was used to generate the distances, linkages, and the final dendrogram.

Robustness coefficients were used to rank-order knockout (overexpression) experiments in terms of the greatest unique responses and identify species which were linearly distinguishable. The response of the knockout (overexpression) was measured in terms of the robustness coefficients. The LRC had desirable linear properties, such that no response (no change in trajectories from wild-type) returns a value of zero and similar negative and positive responses have different directions but similar magnitudes. We considered the unique component of the response to be the orthogonal component in LRC space and the magnitude of the response to be the Euclidean norm. The orthogonal components and there magnitude were identified for each parameter set in the ensemble by first choosing the knockout (overexpression) with the greatest magnitude, *x*_1_ and placing it in the empty set 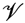. The knockout (overexpression) *x*_1_ defines the orthogonal directions in the LRC space. We then calculated the orthogonal components for all remaining knockouts(overexpressions) relative to *x*_1_, and added the knockout (overexpression) species with the greatest orthogonal magnitude to set 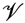. In general the components of all remaining *x_i_* orthogonal to set 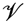 were calculated and the largest was moved into set 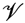;. This process was continued until all knockout (overexpression) species, *x_i_* were added to set 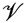. Mathematically two species were considered distinguishable if and only if they were linearly independent (the orthogonal components were non-zero). We considered a threshold value of one or five and performed a student t-test (Matlab Statistical Toolbox, The Mathworks, Natick, MA) to identify which species had orthogonal components above the threshold with a 95% confidence over the ensemble.s

## Author’s contributions

J.V. and A.C. conceived the experiment(s), A.C. conducted the experiment(s) and analyzed the results. A.C. and L.A. wrote the manuscript. All authors reviewed and edited yje manuscript.

## Acknowledgements

We gratefully acknowledge the suggestions from the anonymous reviewers to improve this manuscript.

## Accession codes

Not applicable.

## Supplemental materials

### Molecular basis of the Unfolded Protein Response (UPR)

Protein folding is strategically important to cellular function. Secreted, membrane-bound, and organelle-targeted proteins are typically processed and folded in the endoplasmic reticulum (ER) in eukaryotes^1–3^. Intracellular perturbations caused by a variety of stressors disturb the specialized environment of the ER leading to the accumulation of unfolded proteins^4, 5^. Normally, cells ensure that proteins are correctly folded using a combination of molecular chaperones, foldases, and lectins^1^. However, when proper folding can not be restored, incorrectly folded proteins are targeted to ER Associated Degradation (ERAD) pathways for processing^3^. If unfolded or misfolded proteins continue to accumulate, eukaryotes induce the unfolded protein response (UPR).

UPR is a complex signaling program mediated by three ER transmembrane receptors: activating transcription factor 6 (ATF6), inositol requiring kinase 1 (IRE1) and double-stranded RNA-activated protein kinase (PKR)-like endoplasmic reticulum kinase (PERK). UPR performs three functions: adaptation, alarm, and apoptosis. During adaptation, the UPR tries to reestablish folding homeostasis by inducing the expression of chaperones that enhance protein folding. Simultaneously, translation is globally attenuated to reduce the ER folding load while the degradation of unfolded proteins is increased. If these steps fail, the UPR induces a cellular alarm and apoptosis program. The alarm phase involves several signal transduction events, ultimately leading to the removal of the translational block and the down-regulation of the expression and activity of pro-survival factors such as the B-cell lymphoma 2 (Bcl2) protein. After the alarm phase, cells can undergo apoptosis, although ER stress can also initiate autophagy^6–12^. Thus, ER folding homeostasis strongly influences physiology^5^. Aberrant protein folding and UPR have been implicated in a number of pathologies. For example, the onset of diabetes^74^ as well as myocardial ischaemia, cardiac hypertrophy, atherosclerosis, and heart failure^75^ have all been linked with aberrant folding or UPR signaling.

### The folding cycle, quality control and ER associated degradation (ERAD)

Newly synthesized polypeptide chains enter the ER through a peptide translocon in the ER membrane composed of four proteins, Sec61P (heterortrimeric complex of proteins containing *α,β,γ* subunits) and TRAM^76^. Upon entering the ER, these nascent chains begin to fold, often as they are being co-translationally modified^77^. The folding quality of proteins in the ER is maintained by an in-built quality control (QC) system which ensures proteins are in their native folded state before exiting the ER^4, 78^. A protein is correctly folded if it has attained its native conformation after required co-or post-translational modifications. On the other hand, exposed hydrophobic regions, unpaired cysteine residues, or aggregation are all markers of an unfolded or misfolded conformation^78^, which leads to subsequent retro-translocation to the cytosol. Once in the cytosol, these unfolded or misfolded proteins are degraded by the ubiquitin proteasome system^79^. Hydrophobic unfolded or misfolded queues are recognized in the ER by molecular chaperones which bind these queues and increase the probability of correct folding^80–82^. For example, the HSP70 family of chaperones recognize, in an ATP-dependent manner, exposed hydrophobic patches on a broad spectrum of unfolded or misfolded proteins^3^. Repeated binding and release of HSP70 chaperones ensures that incorrectly folded proteins do not exit the ER^3^. One critical member of the HSP70 family is BiP or GRP78. BiP consists of an N-terminal ATPase domain and a C-terminal peptide binding domain^83^. BiP also regulates the activation of the three transmembrane ER stress transducers: PERK, ATF6, and IRE1. Normally, BiP is bound to these ER receptors, blocking their activation. However, in the presence of exposed hydrophobic residues BiP disassociates, allowing PERK, ATF6, and IRE1 activation. Overexpression of BiP leads to reduced activation of IRE1 and PERK^84, 85^. The PERK and ATF6 branches are thought to be activated before IRE1^34^; this ordering is consistent with the signals that each branch transduces. The PERK and ATF6 pathways largely promote ER adaptation to misfolding, while IRE1 has a dual role, transmitting both survival and pro-apoptotic signals.

### Double-stranded RNA-activated protein kinase (PKR)-like endoplasmic reticulum kinase (PERK) pathway

The PERK branch of UPR transduces both pro-survival as well as pro-apoptotic signals following the accumulation of unfolded or misfolded protein in the ER. PERK is a type I transmembrane protein, composed of a ER luminal stress sensor and a cytosolic protein kinase domain. Dissociation of BiP from the N-terminus of PERK initiates dimerization and autophosphorylation of the kinase domain at T98 1^86^. The eIF2*α* protein, which is composed of three subunits, is critical to translation initiation in eukaryotes, including GTP-dependent start-site recognition^87^. Activated PERK can phosphorylate eIF2*α* at S51^88, 89^, which leads to three downstream effects. First, phosphorylated eIF2*α* globally attenuates translation initiation (Not included in the current model). Decreased translation reduces the influx of protein into the ER, hence diminishing the folding load. Translation attenuation is followed by increased clearance of the accumulated proteins from the ER by ERAD and expression of prosurvival genes. For example, PERK activation induces expression of the cellular inhibitor of apoptosis (cIAP)^38^. Interestingly, decreased protein translation is not universal; genes with internal ribosome entry site (IRES) sequences in the 5′ untranslated regions bypass the eIF2*α* translational block^28^. One of the most well-studied of these, *ATF4*, encodes a cAMP response element-binding transcription factor (C/EBP)^37^. ATF4 drives the expression of pro-survival functions such as amino acid transport and synthesis, redox reactions, and protein secretion^39^. Taken together, these effects seem to be largely pro-survival. However, ATF4 can also induce the expression of pro-apoptotic factors. For example, ATF4 induces the expression of the transcription factor C/EBP homologous protein (CHOP), which is associated with apoptotic cell-death. CHOP (also known as GADD153) is a 29 kDa protein composed of an N-terminal transcriptional activation domain and a C-terminal basic-leucine zipper (bZIP) domain that is normally present at low levels in mammalian cells^40^. The transcriptional activator domain is positively regulated by phosphorylation at S78 and S81 by p38 MAPK family members^90,91^while the bZIP domain plays a key role in the homodimerization of the protein^91, 92^. CHOP activity promotes apoptosis primarily by repression of Bcl2 expression and the sensitization of cells to ER-stress inducing agents^29, 30^.

### Activating transcription factor 6 (ATF6) pathway

ATF6 activation involves a complex series of translocation and irreversible proteolytic processing steps, ultimately leading to the up-regulation of a pro-survival transcriptional program, in the presence of unfolded or misfolded proteins. ATF6 is a 90 kDa ER transmembrane protein with two homologs: ATF6*α*^48,49^and ATF6*β*^93–95^. In the current model, only ATF6*α* is included. Similar to IRE1 and PERK, ER stress leads to the dissociation of BIP from the N-terminus of ATF6, followed by translocation and activation. N-terminal golgi localization sequences (GLS1 and GLS2) seem to be involved with BiP regulation of ATF6. BiP binding to the N-terminal GLS1 promotes the retention of ATF6 in the ER^96^. On the other hand, the GLS2 domain was required to target ATF6 to the golgi body following BiP dissociation from GLS1^96^. Unlike the previous two kinase pathways, ATF6 activation does not involve phosphorylation of a C-terminal kinase domain. Rather, after translocated to the golgi, ATF6 undergoes regulated intramembrane proteolysis (RIP); the luminal domain is first cleaved by serine protease site-1 protease (S1P) followed by metalloprotease site-2 protease (S2P) cleavage^48,97–99^. Cleavage at the juxtamembrane site allows the 50 kDa transcriptional domain of ATF6 to be translocated to the nucleus where it regulates the expression of genes with ATF/cAMP response elements (CREs)^100^ and ER stress response elements (ERSE) in their promoters^52, 101^. Cleaved ATF6 induces a gene expression program, in conjunction with other bZIP transcription factors and required co-regulators, such as nuclear factor Y (NF-Y)^52, 53^, that increases chaperone activity as well as the degradation of unfolded proteins^46, 102^. For example, ATF6 upregulates BiP, protein disulfide isomerase (PDI) and ER degradation-enhancing alpha-mannosidase-like protein 1 (EDEM1) expression. Additionally, ATF6 induces the expression of the X box-binding protein 1 (XBP1) which, after processing by activated IRE1*α*, induces the expression of chaperones. The ATF6-induced gene expression program is also cytoprotective. For example, ATF6 induces regulator of calcineurin 1 (RCAN1) expression^31^. RCAN1 sequesters calcineurin^31^, a calcium activated protein-phosphotase B, that dephosphorylates Bcl2-antagonist of cell death (BAD) at S75 or S99^41^. This leads to sequestering of Bcl2 by Bad, which inhibits its downstream anti-apoptotic activity^41^.

### Inositol-requiring kinase 1 (IRE1) pathway

IRE1 initiates a program with both pro-survival and pro-apoptotic components in the presence of misfolded or unfolded proteins. IRE1 is a 100 kDa type I ER transmembrane protein with both an endoribonuclease and a serine-threonine kinase domain^3^. IRE1 has two homologs, IRE1*α* and IRE1*β*; IRE1*α* is expressed in a variety of tissues^103^ while IRE1*β* is found only in the intestinal epithelia^103, 104^. In the current model only IRE1*α* has been considered. The N-terminus of IRE1, located in the ER lumen, senses unfolded or misfolded proteins through its interaction with BiP^105–107^. Normally BiP is bound to the N-terminus of ire1^84,108,109^. However, in the presence of unfolding queues BiP dissociates and is sequestered by the unfolded or misfolded proteins^110^. Subsequently, IRE1 is activated by homooligomerization followed by autophosphorylation of the C-terminal kinase domain at S724^106,111–113^. IRE1 activation enables both its kinase and endoribonuclease activities to transduce signals simultaneously through two distinct signaling axes. The endoribonuclease activity cleaves a 26-nucleotide intron from the XBP1-mRNa^47,114,115^which generates a 41 kDa frameshift variant (sXBP1) that acts as a potent transcription factor. sXBP1 homodimers, along with co-regulators such as nuclear factor Y (NF-Y), regulate the expression of a variety of ER chaperones and protein degradation related genes^50, 51^. Cytosolic IRE1*α* dimers interact with adaptors such as tumor necrosis factor receptor-associated factor 2 (TRAF2) to drive signal-regulating kinase (ASK1) activation and then subsequently cJUN NH_2_-terminal kinase (JNK) and p38MAPK activation^33^. ASK1 activity is regulated by phosphorylation/de-phosphorylation at several sites as well as by physical interaction with other proteins. ASK1 phosphorylates and activates two downstream kinases, MMK4 and MMK3 which in turn activate JNK and p38 MAP kinase, respectively. JNK is activated by dual phosphorylation at T183 and Y185 by MMK4^116^. Activated JNK activates the proapoptotic Bcl-2 family member Bim by phosphorylation at S65^42, 43^. JNK activation also regulates the activity of anti-apoptotic protein Bcl^44, 117^. Active JNK1 inhibits Bcl2 via phosphorylation at sites T69, S70 and S87^117^. Ultimately, inhibition of Bcl2 and the activation of Bim leads to BAX/BAK dependent apoptosis. Thus, signals initiated from the cytosolic kinase domain of IRE1*α* are largely pro-apoptotic. IRE1*α* activity is regulated by protein serine/threonine phosphatase (PTC2P).

### ER stress-induced apoptosis

Ultimately, if UPR fails to restore ER homeostasis, cells initiate terminal programs such as apoptosis. A common biomarker of apoptosis is the activation of aspartate-specific proteases, collectively known as caspases^118^. Caspases rapidly dismantle cell cycle, cytoskeletal and organelle proteins by proteolytic cleavage. There are two pathways that result in caspase activation in response to apoptotic signals; the death-receptor and the stress mediated pathways. The death-receptor pathway is marked by ligand-mediated activation of death receptors on the plasma membrane. The alternative pathway for caspase activation is mediated by cellular stress e.g., ER stress. Caspases are activated from their zymogens (procaspases), in response to various death cues. First, the initiator caspases, caspase-8 and caspase-9, are activated in response to death cues^119^. This is followed by the activation of executioner caspases, such as caspase-3, caspase-6 and caspase-7. Activated executioner caspases proteolytically process several substrates, facilitating cell death. They also activate initiator caspases, forming a positive feedback loop. Activation of both the PERK and IRE1 pathways modulate stress-induced apoptosis through their regulation of Bcl2 expression and activity. Overall, stress induced apoptosis can occur through both mitochondrial-dependent and independent pathways. Stress signals cause oligomerization of pro-apoptotic proteins, such as Bax and Bak. These proteins are normally sequestered at the mitochondrial outer membrane by the survival protein Bcl2, under non-apoptotic conditions^120^. Once Bax and Bak oligomerize, they insert into the mitochondrial membrane and breach membrane integrity^121^. This results in a net efflux of cytochrome-c from the mitochondria to the cytosol and the initiation of the well-studied Apaf-1 mediated caspase-9 activation pathway. Stress induced mitochondrial-independent apoptotic pathways are not well understood. Currently, caspase 12 has been suggested as a possible ER-stress apoptotic mediator^34,122,123^. However, caspase 12 is not expressed in human. Moreover, there is considerable debate about its role in stress-induced apoptotic cell-death^124^.

## Model Building

### Estimating a population of canonical models using POETs

Using the multiobjective POETs algorithm was used to generate predictive UPR model populations. Each model family was trained and validated on different experimental data. Starting from an initial best-fit initial parameter set (nominal set), more than 25,000 probable models were estimated by POETs from which we selected N = 100 models (25 from each training family) with a Pareto rank of one or less (from approximately 1200 possible choices) for further study. The nominal, training (75 models), and prediction (25 models) errors were calculated for each objective (Table T1). Models used for prediction error calculations for a particular objective were not trained on that objective. The prediction likelihood was statistically significantly better for 31 of the 33 objective functions at a 95% confidence level, compared with random parameter sets generated from the nominal set (Table T1).

Strong Pareto fronts identified in POETs suggested an inability to simultaneously model different aspects of the training data as well as experimental artifacts. Negative feedback was considered to lead to conflicting objectives. For example, XBP1 mRNA measurements (O14) conflicted with CHOP protein measurements (O13), even though these data-sets were taken from the same study and were collected in the same cell-line. XBP1 splicing increased BiP levels, which in turn reduced CHOP protein levels, hence the trade-off. Lastly, in addition to fronts, we also observed strong correlation between objectives. For example, models that performed well for the CHOP protein (O11), also performed well against Procaspase-12 (O22) measurements, even though these were not in the same cell-line or from the same study. Both CHOP and Procaspase-12 are downstream of the IRE1/TRAF2/JNK signaling cascade, so these errors were directly correlated (Fig. 2).

### Signal flow, sensitivity, and robustness analysis of UPR network

Simulated KO and OX studies of key protiens provided insight into the signal flow within the UPR network. Interestingly, PERK and ATF4 KO studies revealed a slower and lower amount of BiP production (~ 50%) as compared to WT. However, ATF6 or IRE1 KO did not affect BiP regulation as compared to WT. This highlighted the dominant role of ATF4 in regulation of BiP, which is consistent with experimental evidence^19^. Regulation of BiP was the critical regulator of spliced XBP1 (XBP1s), which in turn acts as a key marker of progression through different stages of UPR (supplementary materials Fig. S2E). ATF4, cleaved ATF6, and XBP1s act as integrators of the signals coming from all the three branches of UPR and furthermore leads to regulation of BiP, thereby leading to a negative feedback or control of UPR signal. Another interesting note was the regulation of pro-apoptosis phenotype via regulation of Bcl2. PERK and ATF4 KO led to delay in the onset of apoptosis (marked by slower and lower reduction of Bcl2 levels, supplementary materials Fig. S2F). This effect could be attributed to the lack of CHOP mediated branch of Bcl2 regulation. On the other hand, IRE1 and CHOP KO leads to drastic reduction in apoptosis (marked by little or no change of Bcl2 levels, supplementary materials Fig. S2F). CHOP KO implicated the importance of CHOP in the down-regulation of Bcl2. IRE1 KO implicated the critical role of IRE1-TRAF2 mediated route of apoptosis.

A few parameter sets for the sensitivity analysis were diversely selected based on the scatter in the CV values (supplementary materials Fig. S1). Infrastructure parameters e.g. nuclear transport, RNA polymerase or ribosome binding were globally critical, independent of stress (black points, Fig. 6). Additionally, apoptotic species and parameters were also important, both in the presence and absence of UPR (yellow points, Fig. 6). Thus, as expected, components such as RNA polymerase, or caspase activation were globally important irrespective of the folding state of the ER. More interesting, however, were coefficients that shifted above or below the 45°-line in the presence of UPR. These points denote differentially important network components. While the majority of parameters and species became more important in the presence of stress, we found a band of parameters (Fig. 6 Inset) that were differentially important under stressed. For example, the rank-ordering of the sensor and stress-transducer modules clearly increased in the presence of UPR; approximately 172 or 15% of the parameters were significantly more important. These parameters were largely associated with adaptation and processing of unfolded or misfolded proteins, e.g., unfolded protein degradation, cleaved ATF6-induced gene expression, IRE1-TRAF2 mediated apoptosis regulation, and RCAN1 regulation. Likewise, 75 or 12% of the species were significantly more important in UPR compared with normal protein loads (data not shown).

Interestingly, upon knockout of any individual feedback branch like that of ATF4, ATF6 and XBP1s, the system overall remains equally robust. However the sensitivity of the alternate feedback components increases. Overall ~ 54 % of the parameters were differentially less sensitive upon removal of BiP feedback as compared to WT. This brings to light how the presence of BiP feedback makes the system more susceptible/sensitive to perturbations. The specific relevance of ATF4 in trageting BIP feedback was most evident upon KO of ATF4 feedback. We distinctly saw increase in sensitivity of feedback components associated with XBP1s and ATF6 (supplementary materials Fig. S4). Upon ATF6 and XBP1s feedback KO, there wasn’t much change in terms of sensitivity of the system. This further attests the key regulatory effect of ATF4 in mediating the positive BiP feedback which is an essential component of the adaptation phase of UPR. Another interesting observation was that when we completely knockout all the feedback branches of BiP in the adaptation phase, the system overall becomes relatively more robust (supplementary materials Fig. S4). We distinctly saw a major shift of sensitivity of BiP upon removal of positive feedback. KO of ATF6 and XBP1s mediated feedback of BiP was seen to have little effect (as marked by robustness coefficients for BiP, supplementary materials Fig. S7). However, ATF4 mediated feedback KO led to significant amount of reduction in BiP levels (supplementary materials Fig. S7) thereby highlighting the significance of ATF4 in BiP feedback. Upon KO of all branches of BiP feedback, we found overall reductions of BiP levels. However, there were two distinct populations. One with a ~ 10 fold reduction in BiP levels while the other had ~ 1000 fold reduction in BiP levels. These two populations could resemble two distinct operational paradigms within UPR. In the first mode of operation feedback, BiP regulation is really strong resulting in drastic reductions in BiP levels and ultimately a stronger and faster UPR response upon knockout of BiP feedback.

### Structural and parametric uncertainty associated with current version of UPR model

First, the cytosolic kinase domain of PERK can be inhibited by the action of the DNAJ family member P58^*Ipk*^. P58^*Ipk*^ was initially discovered as an inhibitor of the eIF2*α* protein kinase PKR^59^. P58^*Ipk*^, whose expression is induced following ATF6 activation, binds to the cytosolic kinase domain of PERK, inhibiting its activity^60, 61^. Inhibition of PERK kinase activity relieves eIF2*α* phosphorylation, thereby removing the translational block. Interestingly, P58^*Ipk*^ expression occurs several hours after PERK activation and eIF2*α* phosphorylation. Thus, P58^*Ipk*^ induction may mark the end of UPR adaptation, and the beginning of the alarm/apoptosis phase of the response^34^. Second, PERK induces a negative feedback loop, through its downstream effector CHOP, involving the direct de-phosphorylation of eIF2*α*. CHOP induces the expression of GADD34 which, in conjunction with protein phosphatase 1 (PP1), assembles into a phosphatase which dephosphorylates the S51 residue of eIF2*α*^62^. GADD34 is a member of the GADD family of genes which are induced by DNA damage and a variety of other cellular stresses^63^. The GADD34 binding partner in this complex appears to be responsible for PP1α recognition and targeting of the phosphatase complex to the ER. Association between GADD34 and PP1 is encoded by a C-terminal canonical PP1 binding motif, KVRF, while approximately 180 residues, near the N-terminus of GADD34, appear to be responsible for ER localization^64^. Currently, little is known about deactivation of ATF6. Recently, XBP1u, the unspliced form of XBP1, has been implicated as a negative regulator for ATF6^65^. Following, the induction of ER stress, two versions of XBP1 exist: XBP1u and sXBP1^65^. In the recovery phase following ER stress, high levels of XBP1u may play a dual role. First, XBP1u binds sXBP1, promoting complex degradation^66, 67^. Second, XBP1u can bind ATF6*α* rendering it more prone to proteasomal degradation^65^. Taken together, these two steps may slow the transcription of ER chaperones and ERAD components during the recovery phase following ER stress. IRE1*α* activity is regulated by several proteins, including tyrosine phosphatase 1B (PTP-1B), ASK1-interactive protein 1 (AIP1) and members of the Bcl2 protein family. PTP-1B has been implicated in a number of IRE1*α* signaling events. The absence of PTP-1B reduced IRE1*α* dependent JNK activation, XBP1 splicing and EDEM transcription in immortalized and primary mouse embryonic fibroblasts^125^. However, no physical interaction between IRE1*α* and PTP-1B was established. On the other hand, AIP1 physically interacts with both TRAF2 and IRE1*α*, suggesting a model in which AIP1 facilitates IRE1*α* dimerization and activation^126^. The C-terminal period-like domain (PER) of AIF1 binds the N-terminal RING finger domain of TRAF2, followed by ASK1-JNK signaling^127^. Thus, based on these findings, Luo *et al.* postulated that AIF1 may be directly involved in the IRE1*α*-TRAF2 complex and its activation of the ASK1-JNK signaling axis^126^. This hypothesis was validated in AIP1-KO mouse studies; AIP1-knockout mouse embryonic fibroblasts and vascular endothelial cells showed significant reductions in ER-stress induced ASK1-JNK activation that was rescued in AIP1 knock-in cells^126^. IRE1*α* has also been shown to directly interact with Bcl-2 family members Bax and Bak. Hetz *et al.* showed that Bax and Bak complex with the cytosolic domain of IRE1*α* and modulate IRE1*α* signaling^128^. Bax and Bak double knockout mice failed to signal through the IRE1*α* UPR branch following tunicamycin-induced ER stress; however, PERK signaling markers, e.g., eIF2*α* phosphorylation, responded normally^128^. This pro-activation role of Bak and Bax may be modulated by one of the few negative regulators of IRE1*α* activity, Bax inhibitor 1 (BI-1). BI-1 is an anti-apoptotic protein that enhances cell survival following several intrinsic death stimuli^129^. Bailly-Maitre *et al.* were the first to suggest that BI-1 may downregulate IRE1*α* and possibly ATF6 activity^130^. BI-1 deficient mice displayed increased XBP1s and enhanced JNK activity in the liver and kidney, while eIF2*α* phosphorylation remained normal under ER-stress conditions^130^. Lisbona *et al.* later showed that BI-1 directly interacts with the cytosolic domain of IRE1*α*, inhibiting its endoribonuclease activity^131^. Interestingly, BI-1 interacts with several members of the Bcl2 protein family e.g., Bcl2 and Bcl-X_L_, even though it has no homology^129^. Members of the HSP family of proteins have also been shown to regulate IRE1*α*. For example, HSP90 interacts with the cytosolic domain of IRE1*α*, potentially protecting it from degradation by the proteasome^68^. HSP72 interaction with the cytosolic IRE1*α* domain has also recently been shown to enhance IRE1*α* endoribonuclease activity^69^. Taken together, these modes of IRE1*α* regulation with the exception of B1-1, largely promote or enhance IRE1*α* signaling. Given the importance of CHOP in regulation of Bcl2, it is vital to establish the exact connectivity. However, while CHOP expression is negatively correlated with Bcl2 levels, there is no CHOP binding site in the *bcl2* promoter^30^. McCullough *et al.* have suggested that the bZIP domain of CHOP could act with other bZIP transcription factors to regulate *bcl2* expression^30^. Thus, it’s likely that the connection between CHOP expression and apoptosis is more complex than simple down-regulation of Bcl2 expression. These missing structural connections shall allow us to establish a detailed model and extract more relevant insights into manipulating UPR.

**Figure S1.**
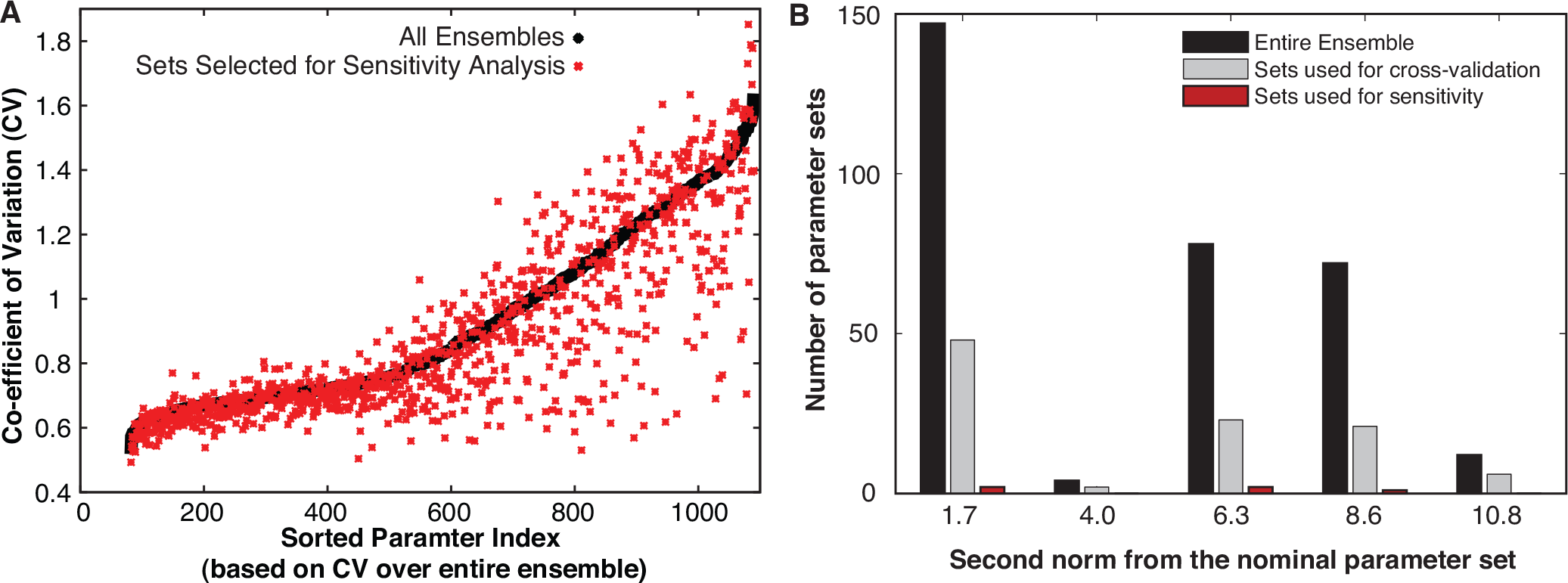
POETs generated an ensemble of models that predicted approximately 94% of the objective functions with a significantly higher likelihood than a random control. (A) The coefficient of variation (CV) for the model parameters ranged from 0.5 - 1.6, where approximately 65% of the parameters were constrained with a CV ≤ 1.0 (black dots). (B) We selected five parameter sets (red dots in A) for further analysis based on CV and distance from the nominal parameter set (based on second norm).

**Figure S2.**
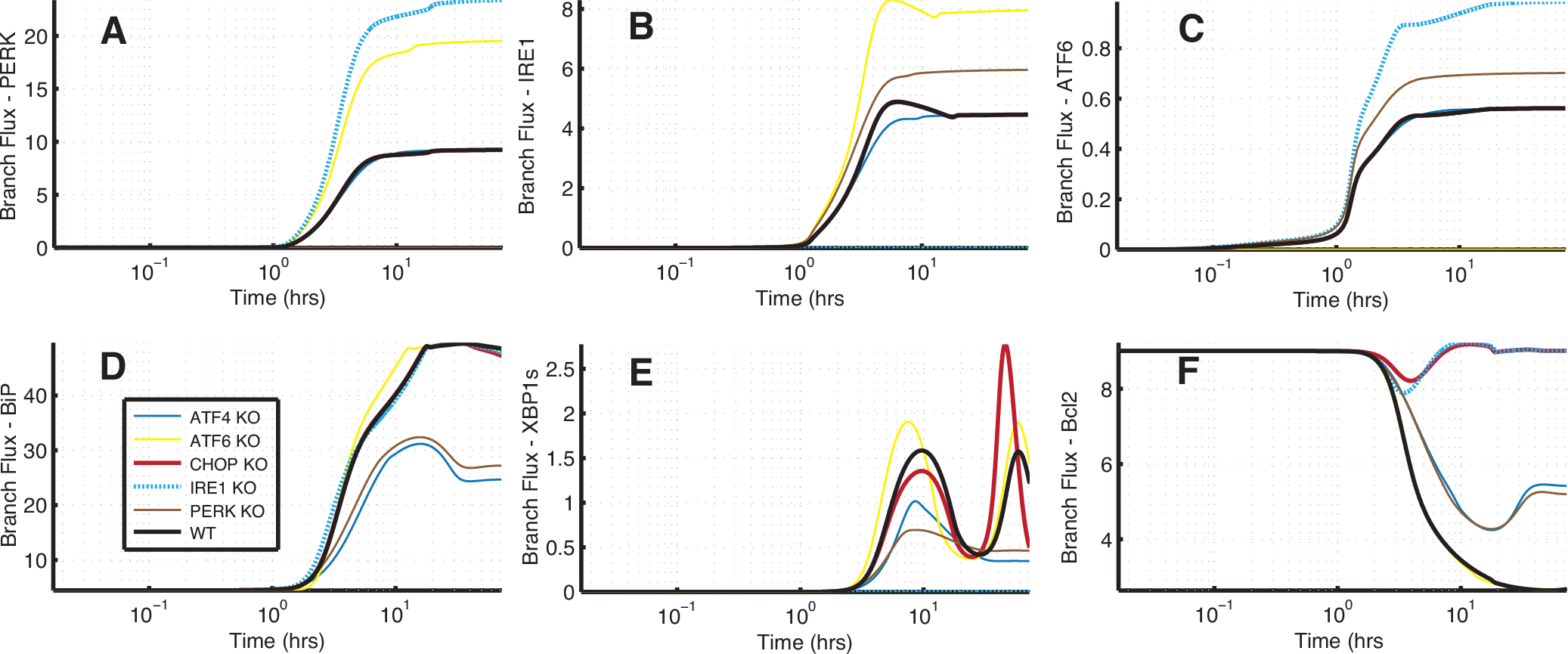
Signal flow analysis using simulated knockout (KO) of key proteins on the UPR system: Simulation results suggest that the three branches in UPR fire simultaneously with varying rates and the state of the cell in terms of adaptation, alarm or apoptosis is a result of counteracting effects of these three prongs of UPR signaling. (A-C) The counteracting effects is seen when knockout of one ER stress transducer leads to enhancement of the other branches of UPR. (D) ATF4, cleaved ATF6 and XBP1s act as integrators of the signals coming from all the three branches of UPR and furthermore leads to regulation of BiP, thereby leading to a negative feedback or control of UPR signal. PERK and ATF4 KO studies revealed a slower and lower amount of BiP production (~ 50%) as compared to WT. However, ATF6 or IRE1 KO did not affect BiP regulation as compared to WT. (E) Regulation of BiP was the critical regulator of spliced XBP1 (XBP1s), which in turn acts as a key marker of progression through different stages of UPR. (F) PERK and ATF4 KO lead to delay in the onset of apoptosis (marked by slower and lower reduction of Bcl2 levels. This effect could be attributed to the lack of CHOP mediated branch of Bcl2 regulation. On the other hand, IRE1 and CHOP KO leads to drastic reduction in apoptosis (marked by little or no change of Bcl2 levels). CHOP KO, implicated the importance of CHOP in the down-regulation of Bcl2. IRE1 KO implicated the critical role of IRE1-TRAF2 mediated route of apoptosis. Overall flux analysis highlighted the extensive amount of crosstalk within the three branches of the UPR network.

**Figure S3.**
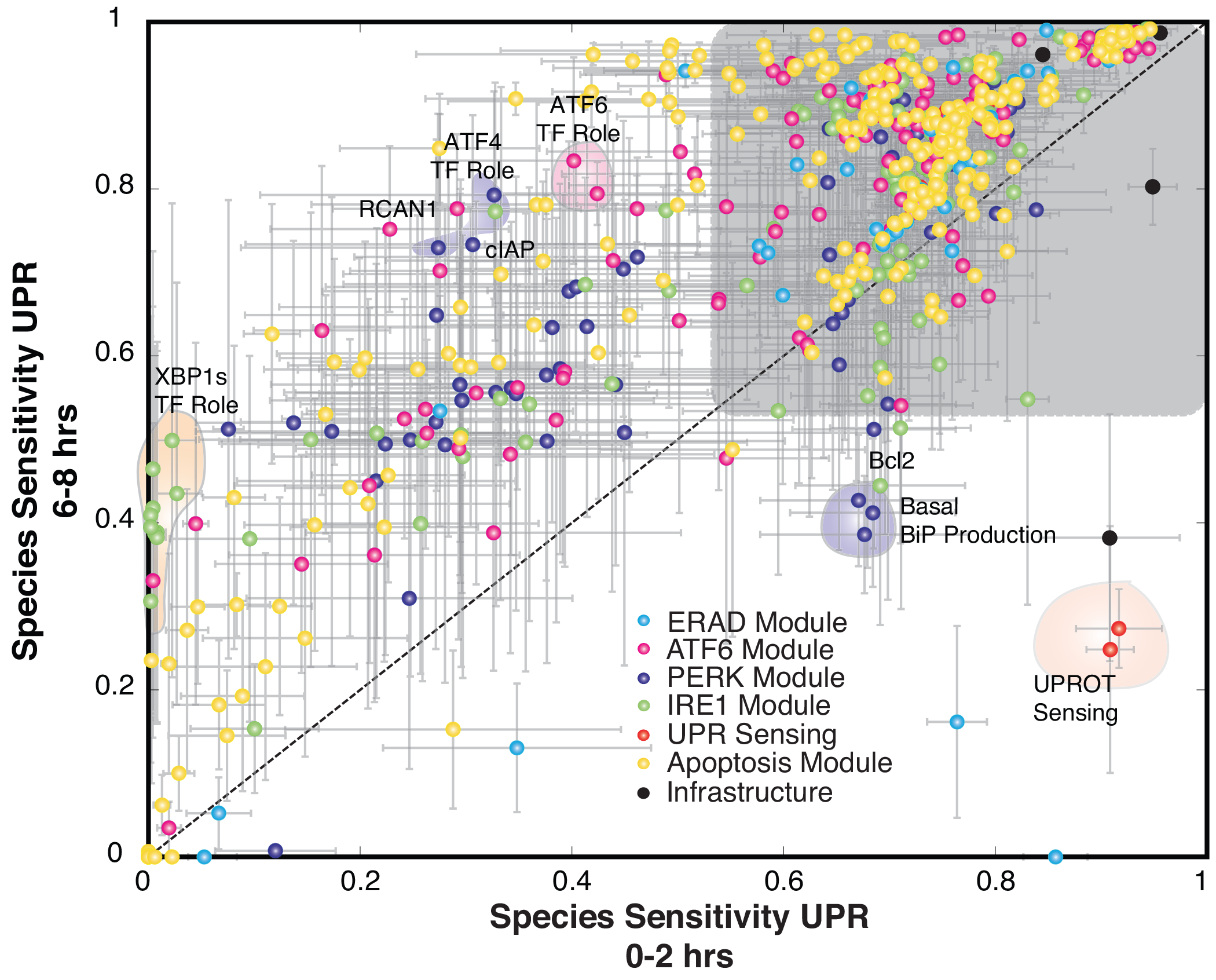
Plot of species sensitivity at earlier (0-2 hrs) versus later (6-8 hrs) time points: Sensitivity analysis was conducted over discrete two hour time windows thereby revealing the time evolution of the importance of UPR network modules. We found that signal integration via the transcriptional activity of ATF6, ATF4 and XBP1s along with RCAN1 and cIAP role in apoptosis were significantly more important at 6-8 hrs as compared to 0-2 hrs time window. This is consistent with the dominant role of the negative feedback via the transcriptional regulation of BiP in UPR. Interestingly, the majority of species rankings were similar as seen in the cluster in the grey box.

**Figure S4.**
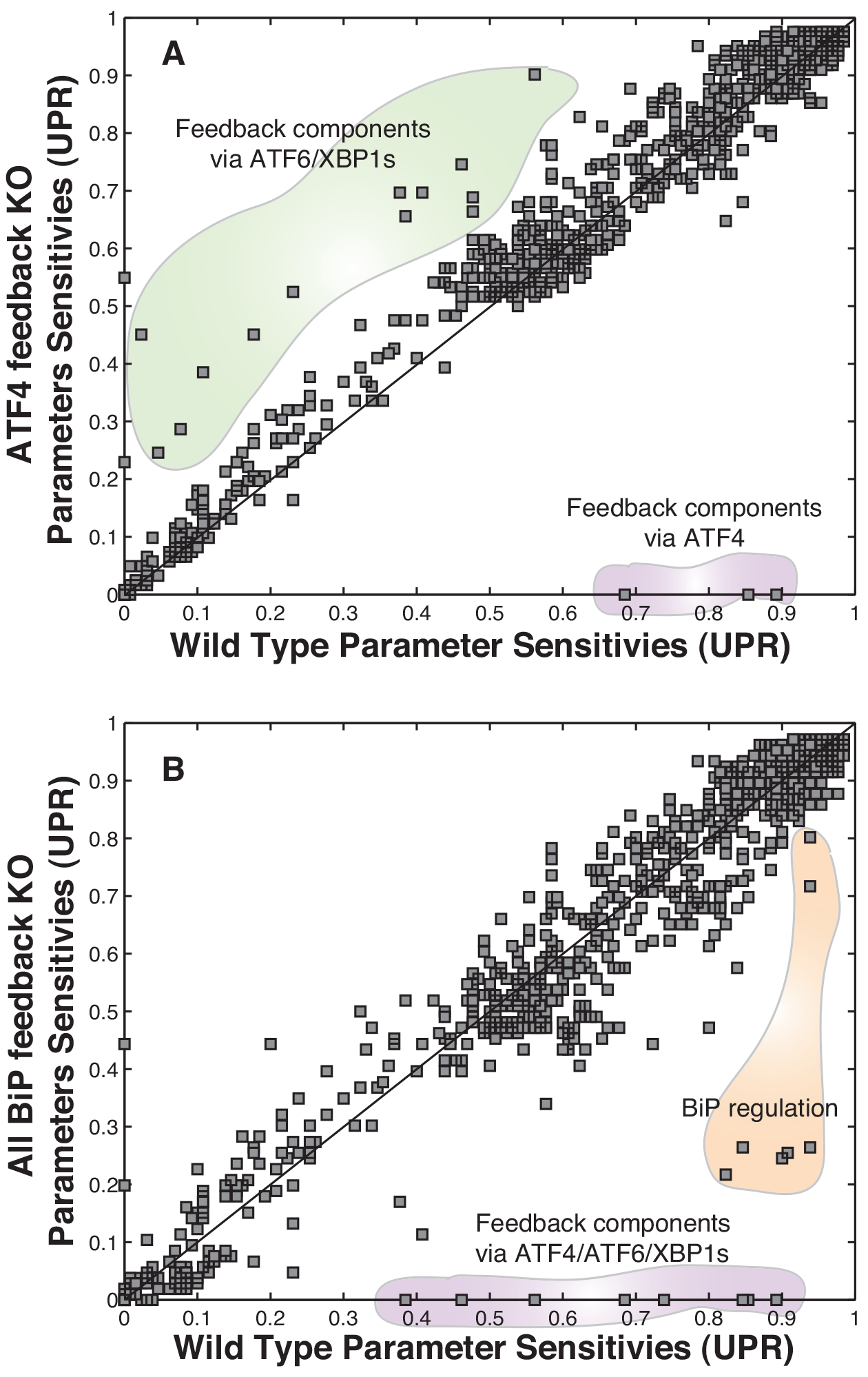
Plot of parameter sensitivity with APAF-1 feedback KO and all BiP feedback KO: Upon knockout of any individual feedback branch like that of ATF4, ATF6 and XBP1s, the system overall remains equally robust. However the sensitivity of the alternate feedback components increases. This was most evident upon ATF4 feedback KO. (A) We saw increase in sensitivity of feedback components associated with XBP1s and ATF6. Upon ATF6 and XBP1s feedback KO, there wasn’t much change in terms of sensitivity of the system (data not shown). This further attests the key regulatory effect of ATF4 in mediating the positive BiP feedback which is an essential component of the adaptation phase of UPR. (B) When we completely knockout all the feedback branches of BiP in the adaptation phase, the system overall becomes relatively more robust. We distinctly saw a major shift of sensitivity of BiP upon removal of positive feedback. Overall ~ 54 % of the parameters were differentially less sensitive upon removal of BiP feedback as compared to WT. This brings to light how the presence of BiP feedback makes the system more susceptible/sensitive to perturbations.

**Figure S5.**
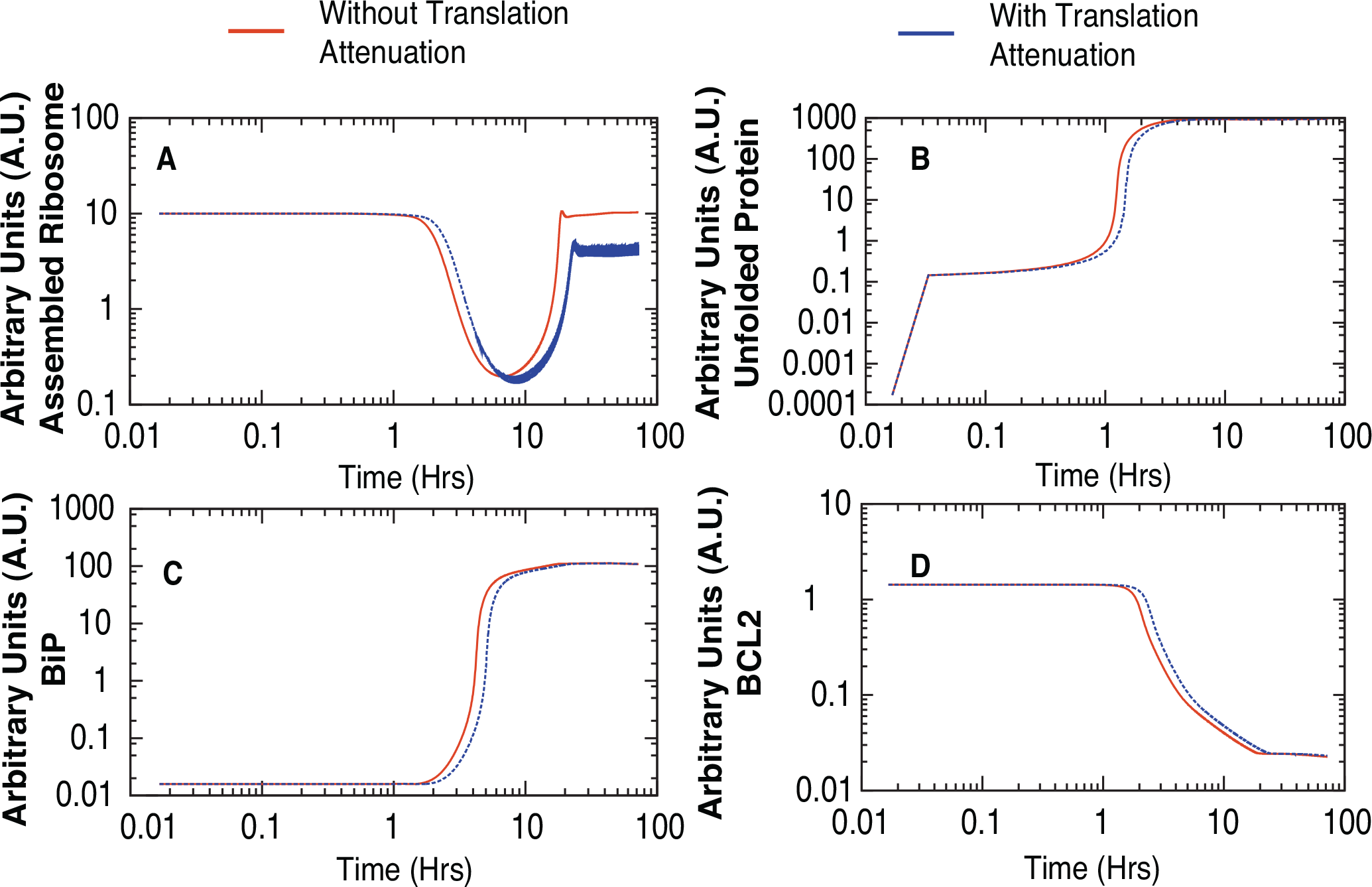
Simulations with translation attenuation built in the model: One of the key aspects which was not included in the current model was translation attenuation. So we simulated that to identify that there isnt much of a change overall in the system except for the tad bit delay in the onset of the responses.

**Figure S6.**
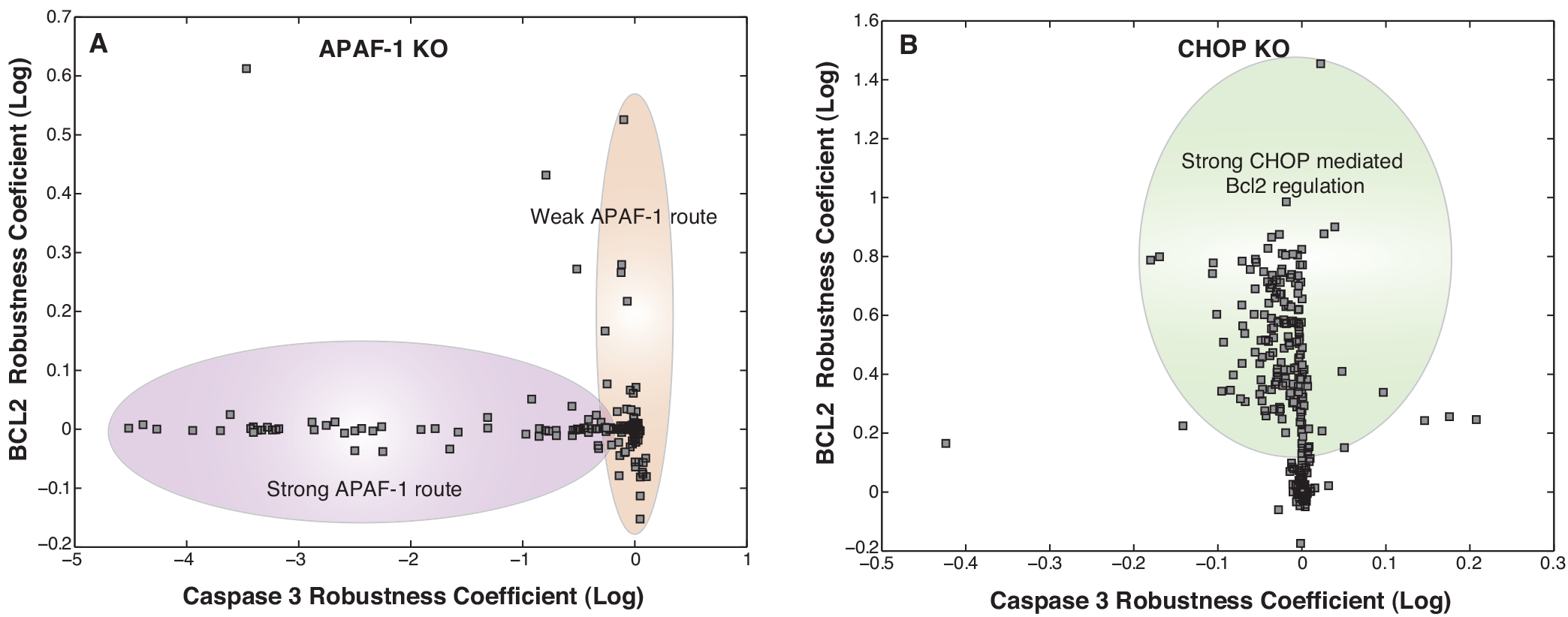
Survival-death phenotypic plane for APAF-1 and CHOP KOs over the entire ensemble: (A) With APAF-1 KO, we found that there were two populations of cells in the ensemble: population 1 where APAF-1 was the dominant regulator of cell-death (marked by enhanced reduction in caspase 3 upon APAF-1 KO) and population 2 where APAF-1 is not the most dominant regulator (marked by reduced effect on Caspase 3 upon APAF-1 KO). (B) Upon CHOP KO, we identified two distinct populations within the ensembles. One with a strong effect of CHOP mediated down-regulation of Bcl2 (marked by ~ 10 fold increase in Bcl2 levels) and the other with very little effect of CHOP on Bcl2 levels. This behavior could be attributed to other conflicting means of regulation of Bcl2 levels.

**Figure S7.**
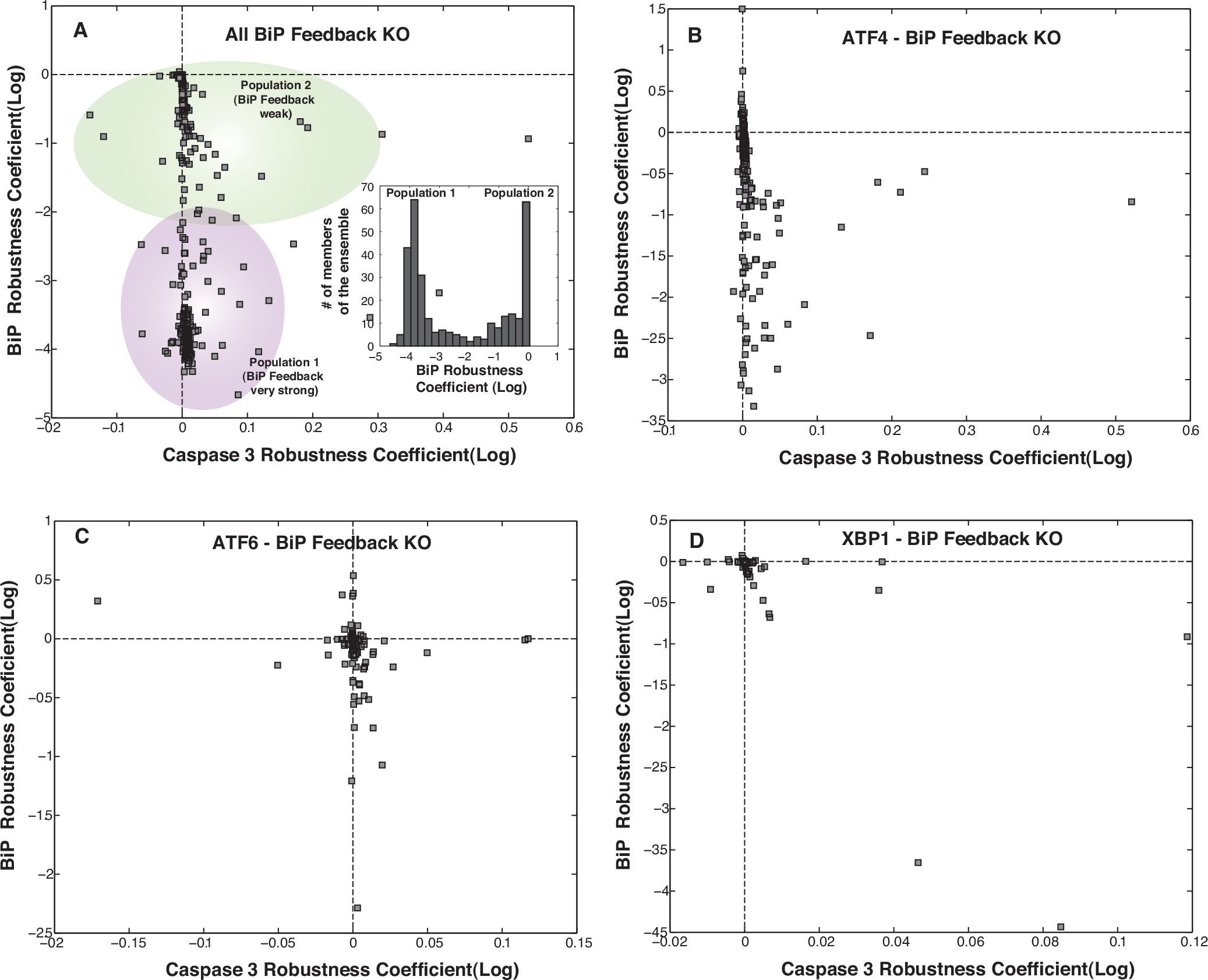
To further investigate the implications of the feedback regulation of BiP via ATF4/ATF6/XBP1s, we simulated KOs of these components over the entire ensemble. (A) Upon KO of all branches of BiP feedback, we found overall reductions of BiP levels. However, there were two distinct sub-populations. One with a ~ 10 fold reduction in BiP levels while the other had ~ 1000 fold reduction in BiP levels. These two populations could resemble two distinct operational paradigms within UPR. In the first mode of operation feedback regulation of BiP is really strong so when we knockout BiP feedback we have drastic reductions in BiP levels and ultimately a stronger and faster UPR response. (B) ATF4 mediated feedback KO led to significant amount of reduction in BiP levels thereby highlighting the significance of ATF4 in BiP feedback. (C-D) However, KO of ATF6 and XBP1s mediated feedback of BiP was seen to have little effect (as marked by robustness coefficients for BiP).

**Table T1.**
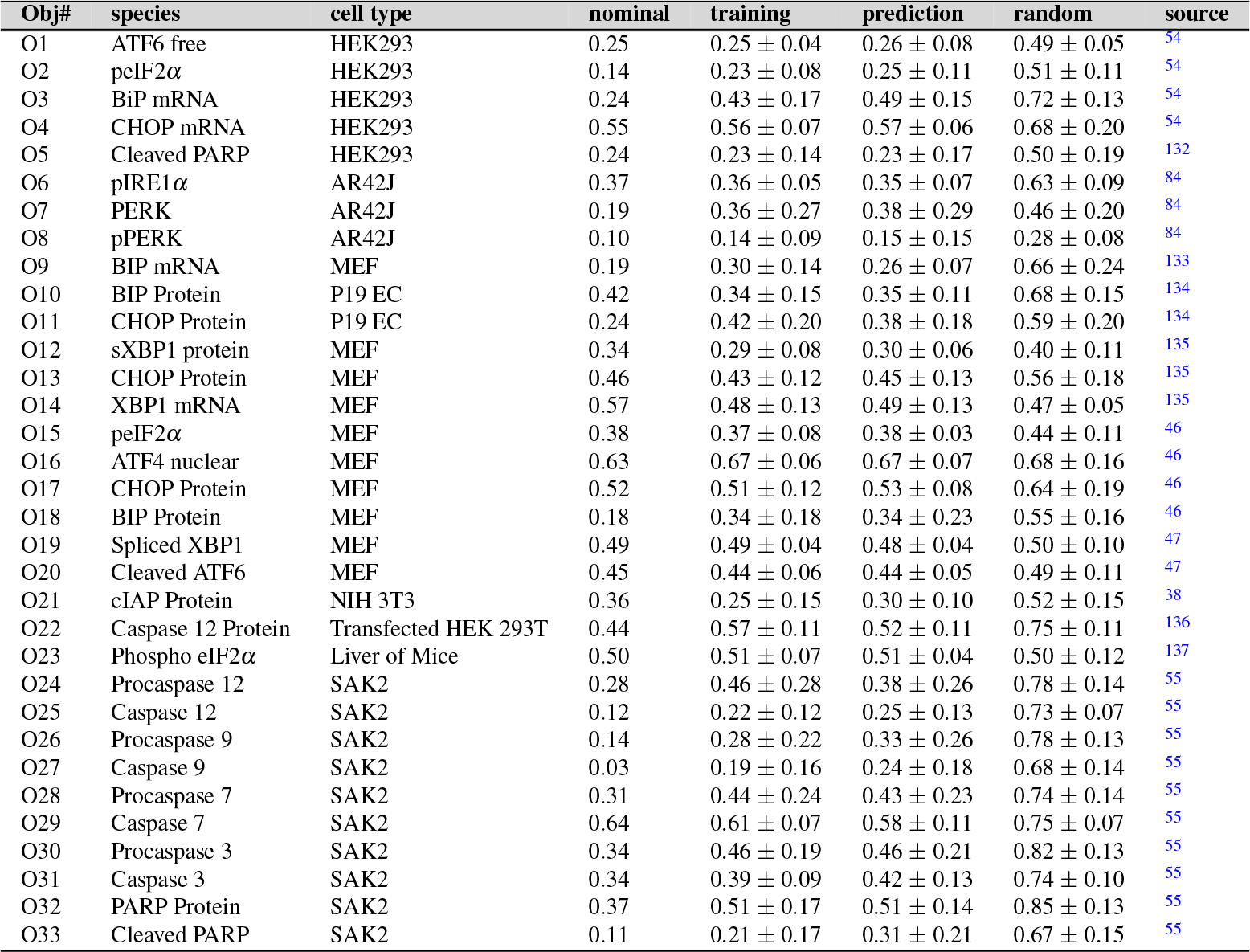
Objective function list along with species, cell-type, nominal error, training error, prediction error, random error with a randomly generated parameter set and the corresponding literature reference.

**Table T2.** Phenotypic response of simulated Gene knockout/overexpression. (**G.O.S** - Gene Overexpression Studies, **G.K.S** - Gene Knockout Studies)

| Increased marker levels | Phenotype | G.O.S (% of cases) | Example genes | G.K.S (% of cases) | Example genes |
| --- | --- | --- | --- | --- | --- |
| BCL2 | Cell Survival | 16 | PERK, ATF4, cIAP, Bcl2, ATF6, RCAN1, Bad, UB | 24 | HSP40, BiP, PERK, eIF2α, ATF4, NFY, JNK, Bad, CHOP, CREB, Procaspase 8 |
| CHOP | – | 5 | eIF2α, CHOP | 3.5 | CEBP, CREB |
| Caspase 3 | Cell Death | 9 | Procaspase 9/8/3/6/7 | 0 | – |
| XBP1 | – | 12 | BiP, cIAP, XBP1, RCAN1 | 3.5 | NFY, TRAF2 |
| BiP | – | 12 | PERK, IRE1, NFY, ATF6, CEBP, CREB, UB | 12 | HSP40, BiP, eIF2α, ATF4, NFY, S1P, S2P |
| PARP | DNA damage | 3.5 | Procaspase 9, PARP | 0 | – |
| ATF4 | – | 20 | HSP40, PERK, BiP, eIF2α, ATF4, IRE1, BAK, TRAF2, ASK1, JNK, p38MAPK, Cal-cinuerin, CHOP, CREB | 5 | NFY, CEBP, CREB |

